# BayesCNet: Bayesian inference for cell type-specific regulatory networks leveraging cell type hierarchy in single-cell data

**DOI:** 10.1101/2025.09.11.675686

**Authors:** Fengdi Zhao, Arkaprava Roy, Weijia Jin, Leeana Peters, Todd Brusko, Qing Lu, Karyn Esser, Ramon Sun, Li Chen

## Abstract

Understanding gene regulatory networks (GRNs) is essential for deciphering biological processes and disease mechanisms. Single-cell multiome technologies now enable joint profiling of chromatin accessibility and gene expression, offering an powerful means to infer cell type-specific GRNs. However, existing methods analyze each cell type independently or aggregate data into pseudo-bulk profiles, limiting their ability to resolve rare populations and capture cellular heterogeneity. We introduce BayesCNet, a Bayesian hierarchical model that jointly infers enhancer-gene linkages across all cell types while leveraging their hierarchical relationships for information sharing. Through extensive simulations, BayesCNet consistently outperforms state-of-the-art methods, with the largest improvements in rare cell types. When applied to real datasets, BayesCNet identifies enhancer-gene linkages with higher accuracy validated by promoter-capture Hi-C data, and reconstructs cell type–specific GRNs that highlight key regulators, demonstrating its power to resolve gene regulatory programs across diverse cell types.

## Introduction

Regulatory DNA elements, such as cis-regulatory elements (CREs) (e.g. enhancer and promoters), are central to the control of gene expression and play critical roles in development and disease^1-3^. They are also central to interpret genome-wide association study (GWAS) signals, the majority of which lie in noncoding regions and may disrupt CREs to alter gene expressions^4^. However, linking CREs to their target genes remains challenging. Enhancer-gene (EG) regulation often occur through chromatin looping, which brings distal regulatory elements into physical proximity with promoters to regulate transcription. Recent sequencing technologies have enables genome-wide profiling of three-dimensional chromatin interactions, such as high-throughput chromosome conformation capture (Hi-C)^5^, Chromatin Interaction Analysis with Paired-End-Tag sequencing (ChIA-PET)^6^ and High-throughput Chromatin Immunoprecipitation combined with Chromosome Conformation Capture (HiChIP)^7^. The resulting chromatin loops, in which one anchor near enhancers and the other anchor near gene promoters, represent gold-standard evidence for EG linkages. Nevertheless, these assays are costly, labor-intensive and typically limited to a few bulk tissues or cell lines. Moreover, their resolution ranges from several kilobases to megabases, which complicates pinpointing precise EG linkage when multiple enhancers or genes reside in the same contact domain. As a result, these assays, while powerful, cannot systematically uncover cell type–specific EG linkages across diverse cellular contexts. Computational approach has therefore been developed to leverage transcriptomic and epigenomic data for inferring EG linkages. Histon modification profiles (e.g. H3K27ac, H3K4me3, H3K4me1) and chromatin accessibility from ATAC-seq can be used to define enhancers and their activity, and correlations between enhancer activity and gene expression, or supervised machine learning approaches^8^ can predict potential regulatory connections. However, this strategy requires large-scale co-profiling RNA-seq data, ATAC-seq, and ChIP-seq data, which is not always feasible. Furthermore, as most existing studies are based on bulk tissues, they overlook cell type-specific EG linkages spanning multiple distinct cell types or cellular states within a heterogeneous sample.

Recent advances in single-cell technologies now make it possible to profile RNA expression (scRNA-seq), chromatin accessibility (scATAC-seq), and their joint measurement (scMultiome) at single-cell resolution across distinct cell types. These advances open new opportunities to interrogate cell type-specific gene regulatory programs, including EG linkages. A growing number of computational methods have been proposed for inferring EG linkages by leveraging single-cell data. Unsupervised approaches such as Cicero^9^ and ArchR^10^ address sparsity and improve statistical robustness by aggregating similar cells. Cicero constructs metacells using a K-nearest neighbor (KNN) graph and applies graphical lasso to estimate partial correlations among co-accessible DNA elements, from which EG linkages are inferred. ArchR uses a similar approach by generating sample-aware pseudo-bulk replicates and computing Pearson correlations between gene expression and chromatin accessibility across cell aggregates. In contrast, supervised methods infer EG linkages within a predictive modeling framework, treating gene expression as the response and chromatin accessibility at ATAC peaks as predictors, with linkage confidence quantified by regression coefficients or feature importance. For example, DIRECT-NET^11^, employs an XGBoost model to capture nonlinear relationships between chromatin accessibility and gene expression in metacells, inferring EG linkages based on feature importance. SnapATAC^12^ adopts logistic regression to assess associations between binarized chromatin accessibility and gene expression, and SCARlink^13^ applies regularized Poisson regression to directly model gene expression counts with ATAC counts as covariates.

Despite the success of these approaches, they are not explicitly designed to identify cell type-specific EG linkages. While they can be adapted for this purpose by analyzing each cell type independently, they face a critical limitation in rare cell types, where small sample size hinders robust inference. One underutilized source of information is the cell type hierarchy, which captures relationships among cell types such as the cell ontology tree (COT) or cell lineage tree (CLT) (**Fig. 1a-b**). Hierarchical cell type relationships have been leveraged by CellWalker2^14^ to improve cell type annotation of scATAC-seq but have not yet been incorporated into EG linkage inference. To fill this gap, we propose BayesCNet, a Bayesian hierarchical framework, for inferring cell type-specific EG linkage and gene regulatory networks (GRNs). BayesCNet jointly models all cell types within a cell type hierarchy, allowing statistical strength to be shared across related lineages. This design improves inference for cell type– specific EG linkage, particularly in rare cell types with limited data. Importantly, BayesCNet is applicable to multiple data modalities, including scMultiome data with co-assay RNA and ATAC in the same cells or single-cell multimodal (scMultimodal) data with RNA and ATAC profiled in different cells, or scATAC-seq data alone. Through comprehensive simulations, we demonstrate that BayesCNet outperforms state-of-the-art methods, with the most evident improvements observed in rare cell types. In real data applications, BayesCNet-inferred EG linkages showed higher concordance with orthogonal promoter-centered Hi-C evidence in two representative datasets, Peripheral Blood Mononuclear Cells (PBMCs) and hematopoietic differentiation, which exemplify both cell ontology tree and cell lineage tree, as well as scMultiome and scMutimodal data, respectively. To illustrate BayesCNet’s utility in complex traits and diseases, we apply BayesCNet to GWAS summary statistics for Systemic lupus erythematosus (SLE), an autoimmune disease, and platelet count and volume, two heritable hematopoietic traits. BayesCNet-identified cell type–specific gene-linked enhancers showed significantly higher heritability enrichment compared to competing approaches. Building on these EG linkages, BayesCNet further constructed cell type–specific GRNs, identifying transcription factor (TF) hubs with high degree centrality. These hubs revealed both cell type– specific regulators and shared regulators, providing insight into key transcriptional programs and cooperative TF modules that drive cell type–specific gene expression.

**Fig. 1.**
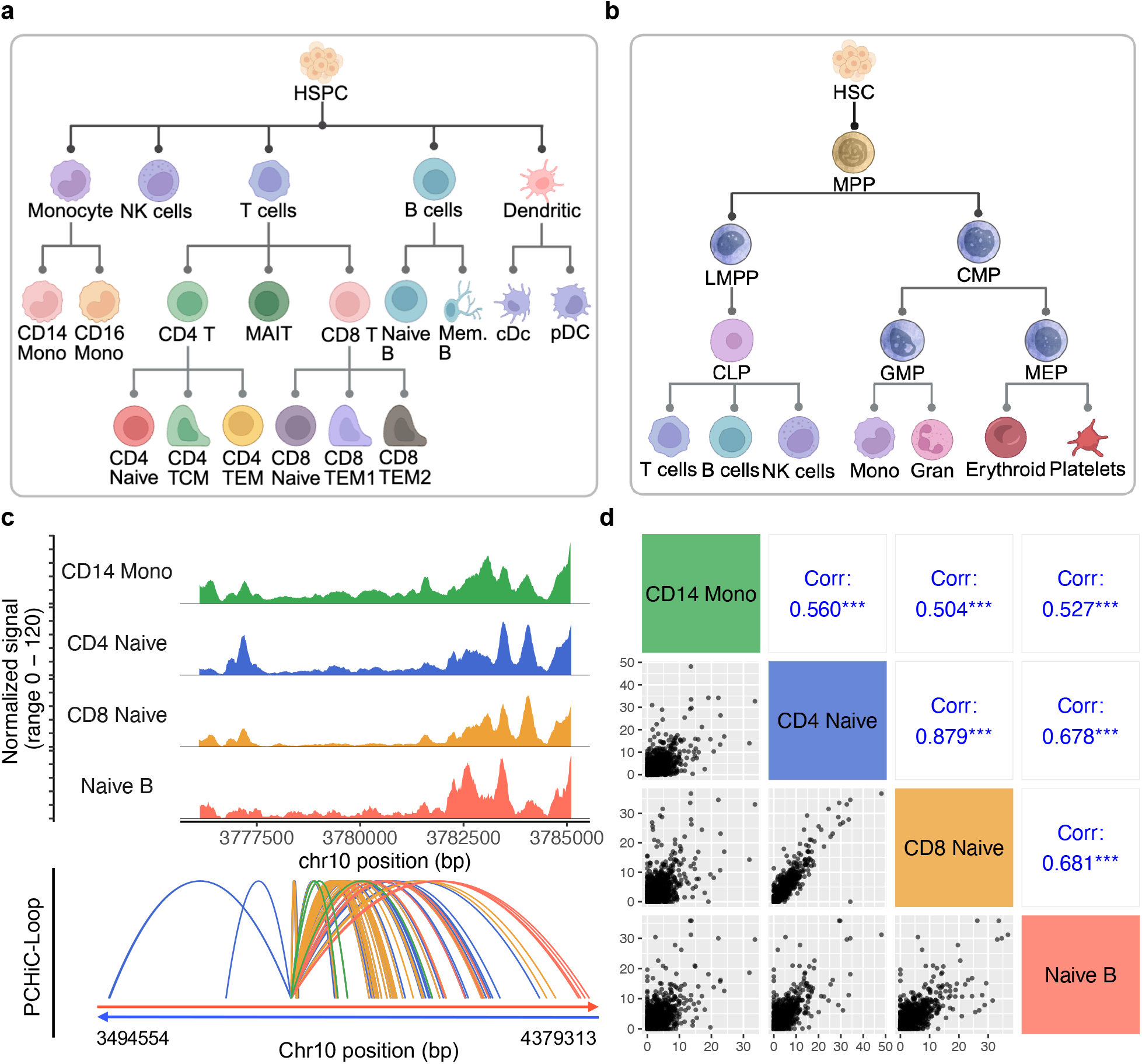
Cell type hierarchy and cell type-specific gene regulation. **a**. Cell ontology tree (COT) of PBMCs, representing a hierarchical classification of cell types. **b**. Cell lineage tree (CLT) of hematopoietic cells (HEMA), reflecting differentiation trajectories from stem and progenitor populations to terminal cell types. **c**. Chromatin accessibility at the KLF6 locus (chr10: 3,775,996–3,785,209) across four PBMC cell types: CD14 Mono, CD4 Naive, CD8 Naive, and Naive B cells. Cell type-specific PCHi-C loops are shown for the four cell types at the KLF6 locus. **d**. Correlation of chromatin interactions among PBMC cell types. Pairwise Pearson correlations of CHiCAGO scores (≥5) are computed based on 10,000 randomly sampled chromatin loops shared between two cell types. The upper triangle reports correlation coefficients, while the lower triangle displays scatterplots of CHiCAGO scores.

## Results

### Cell type hierarchy and cell-type specific gene regulation

We first consider two commonly used forms of cell type hierarchy that serve as structural priors in BayesCNet. Cell Ontology Tree (COT) represents a hierarchical classification of cell types, where each node corresponds to a cell type and edges represent parent–child relationships between broader categories and more specific subtypes. For example, naïve B cells are classified as a child node of B cells, which serve as the parent node. A comprehensive COT of peripheral blood mononuclear cells (PBMCs) is illustrated in **Fig. 1a**. Within this framework, the multi-layer parent–child hierarchy has been widely used for cell type annotation in single-cell data^14^.

However, the COT does not capture differentiation processes and most cell types in a COT are leaf nodes, regardless of whether they are broad categories or fine-grained subtypes. In contrast, the Cell Lineage Tree (CLT) represents biological differentiation trajectories, capturing developmental transitions from stem cells through progenitor states to terminally differentiated cell types. For example, hematopoietic stem cells (HSCs) give rise to multipotent progenitors (MPPs), which further differentiate into common myeloid progenitors (CMPs) and common lymphoid progenitors (CLPs). CMPs generate innate immune and blood cells such as granulocytes, monocytes, and platelets, while CLPs give rise to adaptive immune cells including B cells, T cells, and NK cells (**Fig. 1b)**. Unlike the COT, the CLT explicitly includes both intermediate progenitors (internal nodes) and terminal cell types (leaf nodes). The essential distinction between the two hierarchies is that the COT reflects hierarchical subtype relationships, while the CLT encodes biological differentiation processes. The COT is composed primarily of labeled leaf nodes that may represent either broad or fine-grained categories, whereas the CLT incorporates internal and left nodes to represent both intermediates and fully differentiated types along each lineage.

BayesCNet is developed based on two key biological assumptions: gene regulation is cell type–specific, and cell types share a baseline regulatory program, with similarity being reflected in their proximity in the cell type hierarchy. To illustrate this, we examined the chromatin accessibility near the *KLF6* locus (chr10:3,775,996–3,785,209), an immune response-associated transcriptional regulator^15^, across four representative immune cell types in PBMCs: CD14+ monocytes, CD4 Naïve T cells, CD8 Naïve T cells, and Naïve B cells from PBMC scMultiome dataset (**Fig. 1c**). Accessibility profiles exhibit higher similarity between CD4 and CD8 Naïve T cells than between any other pair of cell types (**Fig. 1c**), which consistent with their closer proximity in the PBMC COT (**Fig. 1a**). Specifically, CD4 Naïve and CD8 Naïve T cells are separated by the shortest path length of 4, compared to a path length of 5 between CD4 Naïve cells and either CD14 monocytes or Naïve B cells. Moreover, cell type–specific promoter– enhancer interactions from PCHi-C further confirmed distinct regulatory programs across these cell types (**Fig. 1c**). To extend this globally, we collected high-confidence chromatin loops (CHiCAGO score ≥ 5) from PCHi-C for the same four cell types and computed pairwise Pearson correlations of CHiCAGO scores using 10,000 randomly sampled shared loops between two cell types. The positive correlations among all cell types indicate a potential shared regulatory program. Consistent with hierarchical proximity, CD4 and CD8 naïve T cells again showed the strongest correlation (Pearson’s R = 0.871), supporting the hypothesis that regulatory similarity aligns with hierarchical relationships among cell types (**Fig. 1d**).

### Overview of BayesCNet

BayesCNet is a Bayesian hierarchical framework for inferring cell type–specific GRNs using single-cell data. It is designed to address two major challenges in single-cell regulatory inference: the sparsity of single-cell measurements and the limited sample size of rare cell types. Unlike methods that construct a cell type–agnostic GRN, or treat each cell type independently, BayesCNet jointly models all cell types within the cell type hierarchy, allowing statistical strength to be shared across related populations. This design improves inference power, particularly for rare populations. The cell type hierarchy can be obtained from prior biological knowledge (e.g., lineage trees or ontology trees) or estimated directly from single-cell data.

An overview of the BayesCNet framework is shown in **Fig. 2**. BayesCNet accommodates multiple single-cell modalities: (i) multiome data, where RNA and ATAC are co-assayed in the same cells, (ii) multimodal data, where RNA and ATAC are profiled in different cells, and (iii) ATAC-only data. This flexibility allows broad applicability across diverse datasets. Single-cell data are preprocessed, clustered and annotated using Seurat and Signac^16,17^. To mitigate data sparsity, transcriptionally similar cells in each cell type are aggregated into metacells using a KNN graph, which enables robust inference at the cell type level (**Fig. 2a**).

**Fig. 2.**
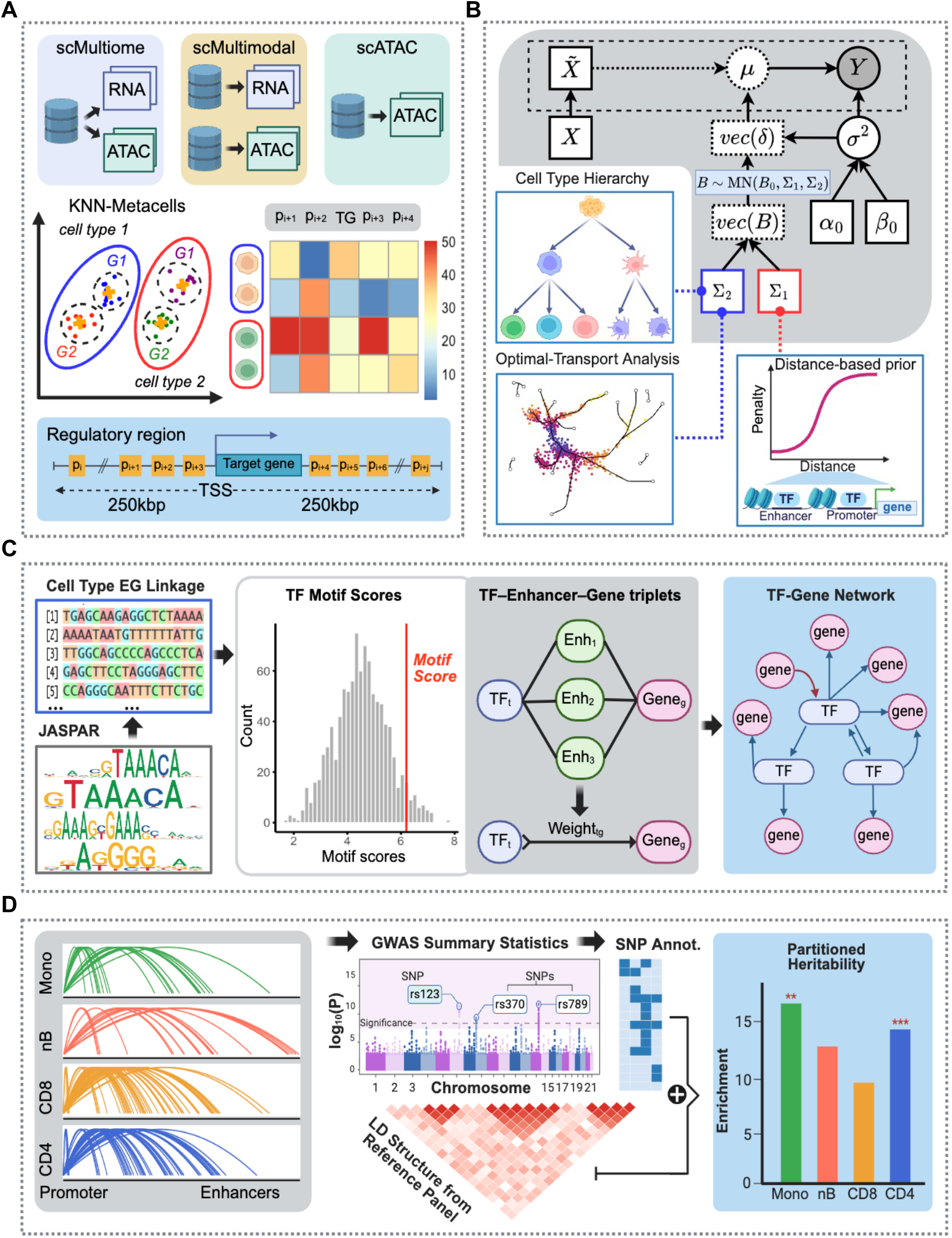
Overview of the BayesCNet. **a**. BayesCNet takes either paired or unpaired single-cell gene expression and chromatin accessibility or chromatin accessibility alone as input. Single cells within each cell type are clustered using a KNN graph and aggregated into metacells. For each gene, BayesCNet models gene expression (or gene activity) as the response variable and chromatin accessibility at peaks within ±250 kb of the transcription starts site (TSS) as covariates. **b**. Cell type–specific enhancer–gene (EG) linkages are inferred using a Bayesian regression framework with structured priors. One prior captures the cell type hierarchy, reflecting developmental lineage or hierarchical cell type relationships, while the other prior encodes genomic proximity and peak co-accessibility. Both priors are represented as covariance matrices. **c**. Predicted cell type–specific EG linkages can be integrated with TF motifs and TF expression to construct cell type–specific TF–gene regulatory networks (TF-GRNs). **d**. Predicted EG linkages can be integrated with GWAS summary statistics to perform partitioned heritability analysis.

These metacells are treated as independent samples in downstream modeling. The BayesCNet framework employs a two-step approach to infer GRNs. First, it aims to infer EG linkages by modeling gene expression as the response variable and chromatin accessibility in ATAC peaks within ±250 kb of the transcription start site (TSS) as the covariates. These ATAC peaks are deemed as putative enhancers^18,19^. When only ATAC data are available, chromatin accessibility within a 500 bp window upstream of the TSS is used as a proxy for gene expression (**Fig. 2a**). Next, BayesCNet incorporates structured priors to guide joint statistical inference of EG linkage for one gene across all cell types. One structured prior captures the cell type hierarchy, reflecting ontological relationships or developmental lineages, while another prior captures peak dependences by modeling genomic proximity and peak co-accessibility. These priors are encoded as covariance matrices that can be derived from prior biological knowledge or estimated empirically from single-cell data (**Fig. 2b**). Second, the resulting cell type–specific EG linkages can be integrated with TF motifs and TF expression to construct TF–GRNs (**Fig. 2c**).

Additionally, EG linkages can be coupled with GWAS summary statistics for partitioned heritability analysis, enabling the identification of trait-relevant regulatory programs (**Fig. 2d**). Full model details are provided in *Methods*.

### BayesCNet improve the inference of enhancer-gene linkages using cell ontology tree

We first evaluated BayesCNet using simulations based on the COT, which organizes cell types according to hierarchical subtype relationships. The goal was to benchmark BayesCNet against competing methods under varying conditions of rare cell type connectivity and sample size. To construct a biologically realistic cell type hierarchy, we adopted a COT resembling that of PBMCs with eight leaf nodes representing terminal cell types (Cell Types 1–8) (**Fig. 3a**). For performance evaluation, we focused on one target cell type under different levels of topological connectivity and sample size. The topological connectivity was defined as the overall strength of a node’s connections to others in the hierarchy, computed using distance-decayed adjacency (*Methods*). A higher value reflects stronger connectivity to other nodes, enabling greater information sharing. Two representative leaf nodes were chosen to represent different connectivity levels: Cell Type 6 (high connectivity = 2.39) and Cell Type 2 (low connectivity=1.91). Sample size was varied by setting the number of metacells for the target cell type to 5 (low), 20 (medium), or 60 (high), while fixing the remaining cell types at the medium level of 20. These values were chosen to match the empirical distribution of metacells observed in PBMC scMultiome dataset (**Supplementary Table 1**). This yielded a 2 × 3 factorial design (connectivity × sample size), resulting in six simulation scenarios, which enabled a systematic evaluation of model performance across both cell type abundance (rare, moderate, abundant) and topological connectivity (low vs. high) (**Fig. 3b**).

**Fig. 3.**
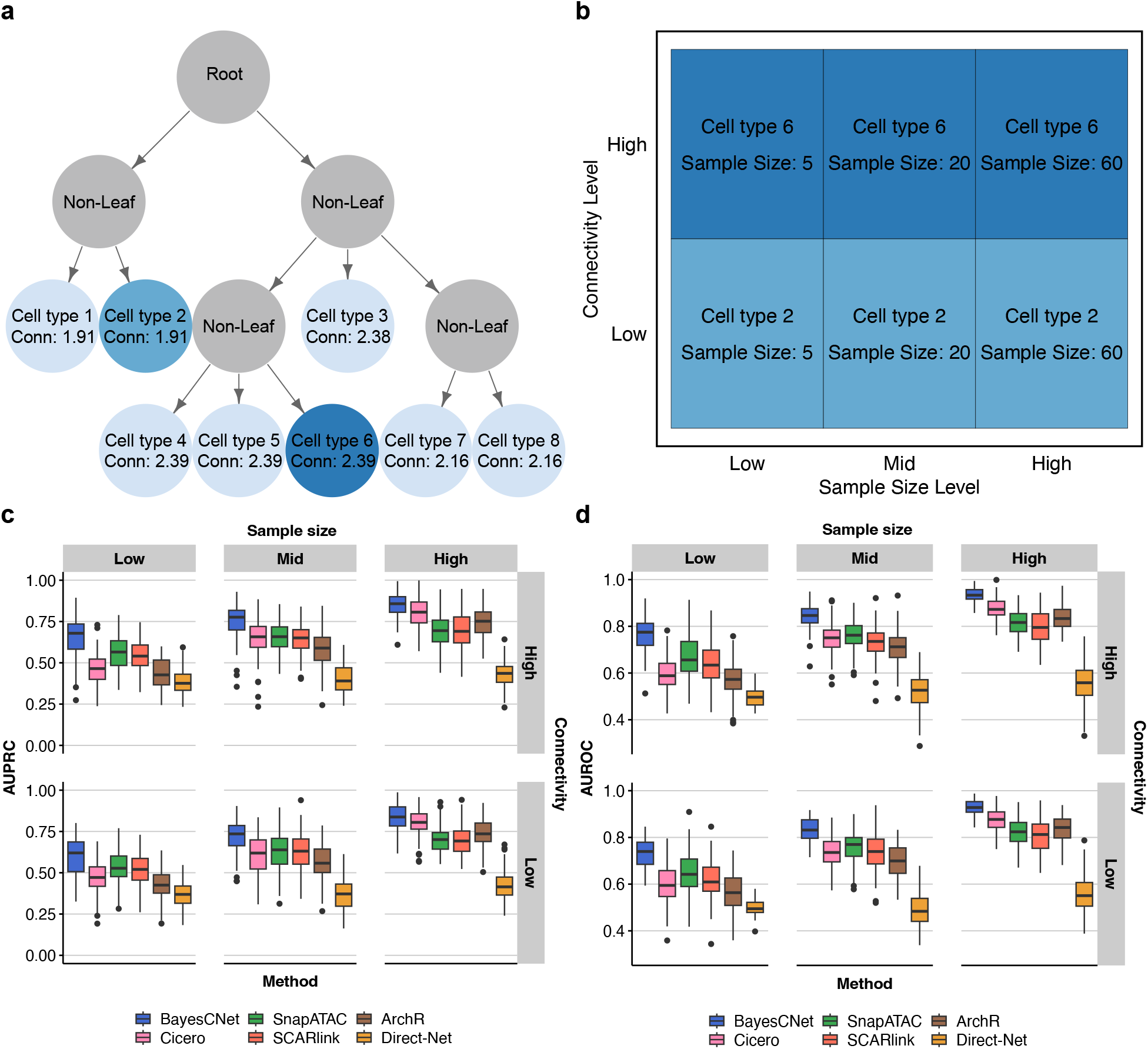
Simulation studies for comparing BayesCNet, which leverages cell type relationship inferred from cell ontology tree (COT), to competing methods for inferring enhancer-gene linkages. **a**. A simulated cell ontology tree, mimicking PBMC cell type hierarchy, for simulation. Two target cell types were selected to represent different connectivity levels: Cell type 2 (low connectivity) and Cell type 6 (high connectivity). Connectivity scores are indicated at each node. **b**. Experimental design of the 2×3 factorial simulation for connectivity and sample size. Each target cell type was simulated at three sample size levels: low (5), medium (20), and high (60) metacells, and two connectivity level: low and high. The sample sizes of other cell types are fixed at the medium level. **c**. AUPRC across 100 simulations per method per scenario. **d**. AUROC across 100 simulations per method per scenario.

Synthetic single-cell RNA and ATAC counts were generated to mimic the statistical and biological properties of PBMC scMultiome dataset. Simulations incorporated both cell type relationships, derived from the PBMC COT, and peak–peak correlations, estimated from empirical chromatin accessibility, to capture realistic dependencies across both cell types and genomic features. Ground-truth EG linkages were embedded through regression coefficients sampled from structured distributions, with large positive values indicating active regulatory links (labeled as 1 if ≥ 0.5 and 0 otherwise). Single-cell ATAC counts were simulated using a Gaussian copula with negative binomial marginals, and single-cell RNA counts were simulated using a log-linear negative binomial model that links accessibility with regression coefficients. To construct metacell data, single cells were aggregated into metacells using K-means clustering. Metacell ATAC counts were obtained by summing accessibility across cells in each cluster, and metacell RNA expression was then simulated using a Gaussian linear model that integrates metacell ATAC counts with regression coefficients. This strategy produced two complementary datasets on both single-cell and metacell resolution, which enabled evaluation of methods tailored to each input type: single-cell–based approaches (e.g., SCARlink, SnapATAC) and metacell-based approaches (e.g., BayesCNet, DIRECT-NET, Cicero, ArchR). Each method produced importance scores for candidate gene–peak pairs, which were compared to ground-truth EG linkages. To ensure robustness, each simulation scenario was repeated 100 times, and performance was summarized using the median AUPRC (mAUPRC) and AUROC (mAUROC). Full details of the data generation procedure are provided in *Methods*.

Across all six scenarios, BayesCNet consistently achieved the best performance, while DIRECT-NET performed the poorest (**Fig. 3c**). In the challenging case of low sample size but high connectivity, representing a rare cell type placed in a highly connected position of the COT, BayesCNet attained the highest mAUPRC (0.679), outperforming the next best supervised methods, SnapATAC (0.565) and SCARlink (0.541). A Wilcoxon rank-sum test confirmed that BayesCNet’s improvement was statistically significant (p= 3.1×10^−9^ vs. SnapATAC). In contrast, ArchR and Cicero, two unsupervised methods, attained mAUPRC values below 0.5, and DIRECT-NET performed worst (0.376). These results highlight BayesCNet’s ability to leverage structured priors from cell type hierarchy, enabling robust inference of EG linkages in rare cell types. The relative performance of competing methods reflected their strategies for handling sparsity. SnapATAC and SCARlink, which operate directly at the single-cell level, preserve granularity and effectively increase the available sample size, partially explaining their stronger performance in low-sample regimes. In contrast, ArchR and Cicero reduce sparsity by aggregating cells into metacells, which stabilizes estimates but reduces effective sample size, leading to weaker performance in low-sample regimes. However, as sample size increased, metacell-based approaches improved substantially and began to outperform single-cell methods, likely due to stronger effect of sparsity reduction by cell aggregation. For example, in the scenario of high-sample and high connectivity, Cicero (mAUPRC = 0.807) and ArchR (0.752) outperformed SnapATAC (0.695) and SCARlink (0.690). Although DIRECT-NET also improved with more data (from 0.376 to 0.436), it remained the weakest method overall, likely reflecting the larger sample size needed to support its more complex nonlinear tree-based model.

BayesCNet maintained strong performance even in the most challenging scenario of low connectivity and low sample size, achieving mAUPRC of 0.620. This remained significantly better than SnapATAC (0.527, p = 1.095×10^−6^) and SCARlink (0.520, p = 2.028×10^−8^), demonstrating the robustness of BayesCNet. Furthermore, BayesCNet benefited from increased connectivity: with sample size fixed low, mAUPRC rose from 0.620 (low connectivity) to 0.679 (high connectivity). This indicates that the position of a rare cell type within the hierarchy affects BayesCNet’s performance. A rare cell type placed at a “hub” node can borrow more information from other cell types and thus improving its performance. Similar connectivity-related gains were observed at medium and high sample sizes, where mAUPRC increased from 0.735 to 0.776 and 0.838 to 0.858. However, this effect diminished as sample sizes grew and sufficient within-cell-type information became available. A parallel trend was observed for sample size effects. With connectivity fixed, mAUPRC steadily increased with sample size, rising from 0.620 to 0.735 to 0.838 under low connectivity and from 0.679 to 0.776 to 0.858 under high connectivity. As expected, competing methods also improved with larger sample sizes. However, for a given sample size, their performance remained largely stable regardless of connectivity.

Performance measured by AUROC followed the same overall trend (**Fig. 3d**). BayesCNet exhibited the strongest advantage in small-sample regimes, with further improvements when higher connectivity increased. Single-cell methods such as SnapATAC and SCARlink benefitted in low-sample scenarios, but their advantage diminished as sample size increased, where metacell-based approaches showed stronger performance. DIRECT-NET consistently performed worst. Overall, these results demonstrate the superior performance of BayesCNet across diverse scenarios, showing robustness to changes in sample size, from rare to abundant cell types, and to variations in topological connectivity, whether cell types are positioned at nodes with high or low connectivity within the hierarchy. Using cell type proportions estimated directly from PBMC scMultiome data, we carried out an additional simulation study and observed that BayesCNet maintained superior performance over competing methods across all cell types (**Supplementary Fig. 1**).

Finally, we benchmarked BayesCNet with an empirical prior (BayesCNet_E) against external methods (**Supplementary Fig. 2**). The empirical prior is calculated using cell embedding based on ATAC or RNA or both modalities (*Methods*). Across all six scenarios, BayesCNet_E consistently outperformed the best competing method, though it remained slightly below BayesCNet with biological priors. For high connectivity, BayesCNet_E achieved mAUPRCs of 0.586, 0.753, and 0.858 for low, medium, and high sample sizes, respectively, compared to SnapATAC (0.565), Cicero (0.657), and Cicero (0.807). For low connectivity, BayesCNet_E attained 0.575, 0.714, and 0.837, again exceeding all competing approaches. A similar trend is observed for mAUROC. These results demonstrate that even without the COT-derived biological prior, BayesCNet with the empirically estimated priors is a strong alternative, further underscoring the robustness and flexibility of the framework.

### BayesCNet improve the inference of enhancer-gene linkages using cell lineage tree

We next evaluated BayesCNet using the second form of cell type hierarchy-CLT. The simulated CLT comprised 10 cell types modeled after hematopoietic differentiation (**Fig. 4a**). Unlike COT, which only classifies terminal subtypes, CLT includes both intermediate and terminal nodes, capturing the continuum of hematopoiesis in which progenitors and differentiated cell types coexist. To assess the impact of connectivity, two representative target cell types are selected: Cell Type 3 (connectivity = 4.74), a highly connected multipotent progenitor, and Cell Type 8 (connectivity = 3.22), a terminally differentiated type with lower connectivity. A key difference in prior specification is that all root, intermediate, and terminal nodes in CLT contribute to the structural prior, whereas only leaf nodes are considered in COT. Sample sizes were assigned based on observed proportions in HEMA scMultimodal dataset (**Supplementary Table 2**), with target cell types receiving 8, 15, or 40 metacells (low, medium, and high), while all other cell types were fixed at the medium level of 15. The simulation followed the same 2 × 3 factorial design as the COT analysis, and data generation proceeded as described above (**Fig. 4b**).

**Fig. 4.**
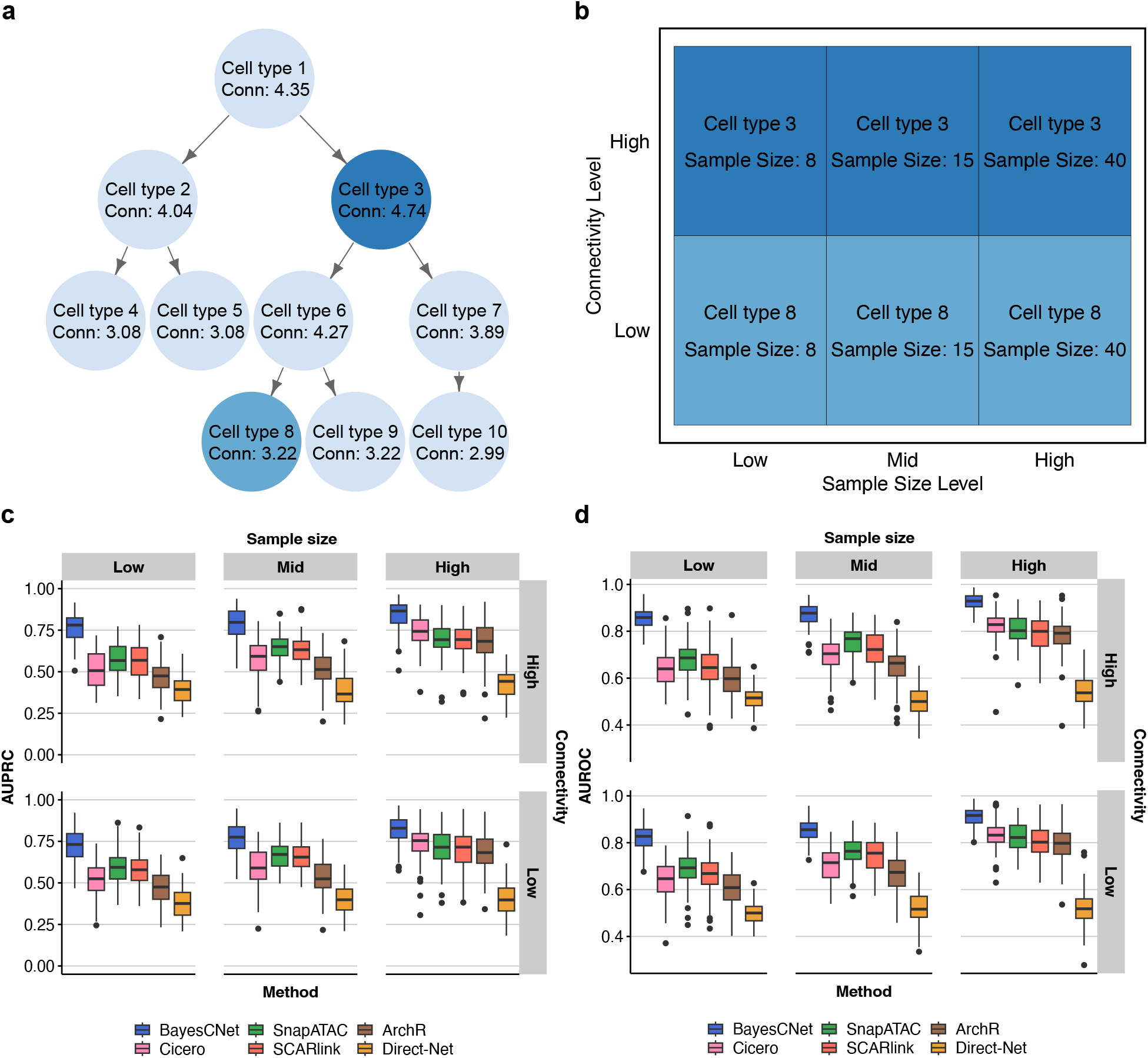
Simulation studies for comparing BayesCNet, which leverages cell type relationship inferred from cell lineage tree (CLT), to competing methods for inferring enhancer-gene linkages. **a**. A simulated cell lineage tree, mimicking hematopoietic differentiation, for simulation. Two target cell types were selected to represent different connectivity levels: Cell type 8 (low connectivity) and Cell type 3 (high connectivity). Connectivity scores are indicated at each node. **b**. Experimental design of the 2×3 factorial simulation for connectivity and sample size. Each target cell type was simulated at three sample size levels: low (8), medium (15), and high (40) metacells, and two connectivity level: low and high. The sample sizes of other cell types are fixed at the medium level. **c**. AUPRC across 100 simulations per method per scenario. **d**. AUROC across 100 simulations per method per scenario.

Across all simulation scenarios, BayesCNet consistently achieved the strongest performance, while DIRECT-NET again performed the worst (**Fig. 4c**). The advantage of BayesCNet was most pronounced under low-sample regimes. For example, in the scenario of high connectivity and low sample size, BayesCNet reached an mAUPRC of 0.781, exceeding SCARlink (0.567) by more than 0.2. Under low connectivity, BayesCNet achieved 0.731, outperforming SnapATAC (0.594) by more than 0.1. These findings show that rare cell types located at hub positions in the lineage tree benefit substantially from BayesCNet’s incorporation of hierarchical priors. As expected, performance of all methods improved with larger sample sizes. Single-cell methods (SnapATAC, SCARlink) performed better than metacell-based methods (Cicero, ArchR) in low-sample settings, while Cicero and ArchR improved markedly at higher sample sizes. DIRECT-NET remained the poorest-performing method throughout. A similar trend was observed when performance was assessed by AUROC (**Fig. 4d**).

Direct comparison of BayesCNet under COT versus CLT revealed stronger gains from CLT priors, particularly at small sample sizes. In the scenario of low-sample high-connectivity, BayesCNet achieved an mAUPRC of 0.781 under CLT, compared to 0.679 under COT. The margin of improvement over the next-best method was much greater under CLT (0.214 vs. SCARlink at 0.567) than under COT (0.114 vs. SnapATAC at 0.565). This advantage arises from the higher connectivity of the target node in CLT (4.74) compared to the most connected node in the COT (2.39). By incorporating intermediate progenitor nodes, CLT yields a higher overall average connectivity than COT, which only includes terminal leaf nodes. A similar benefit was observed in low-sample low-connectivity setting, where BayesCNet reached a mAUPRC of 0.731 under CLT compared to 0.620 under COT. At medium and high sample sizes, BayesCNet performed comparably across CLT and COT (e.g., 0.838 vs. 0.829 for low connectivity at high sample size; 0.858 vs. 0.856 for high connectivity at high sample size). An additional simulation using HEMA scMultimodal-derived cell type proportions confirmed that BayesCNet consistently outperformed competing methods across all cell types (**Supplementary Fig. 3**)

Finally, we assessed BayesCNet_E, which incorporates empirically estimated priors (**Supplementary Fig. 4**). BayesCNet_E consistently outperformed the strongest external methods, though it remained slightly below BayesCNet with biological priors. Under high connectivity, BayesCNet_E achieved mAUPRCs of 0.644, 0.707, and 0.827 for low, medium, and high sample sizes, respectively, compared to SCARlink (0.569), SnapATAC (0.651), and Cicero (0.743). For low connectivity, it attained 0.600, 0.725, and 0.808 across the three sample sizes, again surpassing all external methods. A similar trend was observed when performance was assessed using AUROC (**Fig. 4d**). These findings highlight the flexibility of BayesCNet, showing that even without a biologically defined CLT prior, empirical priors provide a strong and competitive alternative.

### BayesCNet predicts enhancer-gene linkages supported by PCHi-C chromatin interactions

To benchmark BayesCNet on real data, we applied it alongside competing methods to the PBMC scMultiome dataset for inferring EG linkages. BayesCNet followed its standard framework (**Fig. 2a, b**), while competing methods were run using their recommended pipelines and default settings. For metacell-based methods, we set K=50 for KNN graph-based aggregation. Each gene–peak pair was ranked using each method’s own scoring scheme (*Methods*). As an orthogonal gold standard, we used PCHi-C maps of promoter-enhancer interactions across 17 human blood cell types, retaining only high-confidence chromatin loops^20^ (**Supplementary Table S3**). To align PBMC scMultiome dataset with PCHi-C interactions, we directly matched cell types where possible and mapped unmatched cell types by linking child nodes to parent nodes based on PBMC COT (*Methods*). This procedure yielded nine immune cell types with matched PCHi-C loops: CD14 Monocytes, CD16 Monocytes, CD4 Naïve, CD4 TCM, CD4 TEM, CD8 Naïve, CD8 TEM1, CD8 TEM2 and Naïve B. These cell types span a wide range of sample sizes, from abundant CD14 Monocytes to rare Naïve B, providing a natural testbed for assessing performance under different data regimes (**Supplementary Table S4**). For each cell type, we evaluated a fixed set of marker genes (i.e. upregulated protein-coding genes) across all methods. Each method ranked candidate gene–peak pairs per gene using its own scoring scheme. A marker gene was considered evaluable for a given method if at least five of its predicted EG pairs overlapped PCHi-C loop anchors in the matched cell type. For evaluable genes, we computed per-gene AUROC and AUPRC using PCHi-C overlaps as positives and summarized performance across genes with distributions visualized by interquartile range (IQR) (**Fig. 5**).

**Fig. 5.**
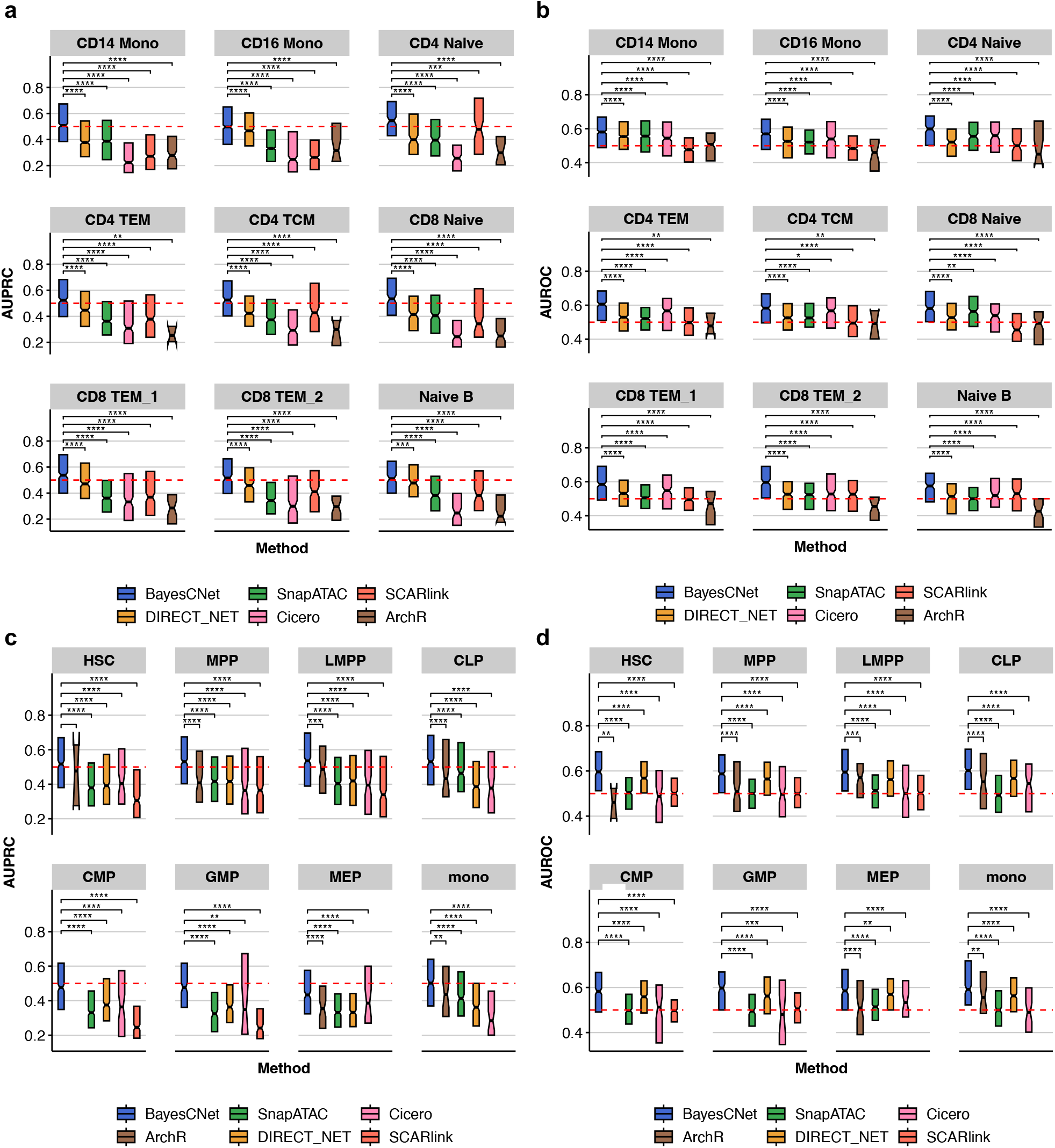
Real data benchmarking using PCHi-C data for predicted enhancer-gene linkages. **a**. AUPRC for cell type-specific gene markers by comparing predicted enhancer–gene linkages and PCHi-C loops across nine immune cell types for PBMC scMultiome dataset **b**. AUROC scores for the same cell types. **c**. AUPRC for cell type-specific gene markers by comparing predicted enhancer–gene linkages and PCHi-C loops across eight progenitor cell types for HEMA scMultimodal dataset **d**. AUROC scores for the same cell types. For all evaluations, only the cell type-specific gene markers with at least five overlapped with PCHi-C loops are considered. Asterisks indicate statistical significance from pairwise Wilcoxon rank-sum tests (p < 0.05; p < 0.01; p < 0.001; p < 0.0001).

On the PBMC scMultiome dataset, BayesCNet consistently ranked first across the nine immune cell types, achieving the highest mAUPRC in all cases (**Fig. 5a**). DIRECT-NET was second best in six cell types, SCARlink in two, and SnapATAC in one. BayesCNet’s advantage was evident in rare populations. For Naïve B (145 cells/3 metacells), BayesCNet reached a mAUPRC of 0.511, surpassing DIRECT-NET (0.478) and substantially outperforming single-cell methods, SnapATAC (0.382) and SCARlink (0.381). An unpaired Wilcoxon rank-sum test confirmed BayesCNet significantly outperformed DIRECT-NET across Naïve B-specific marker genes (p = 3.66 × 10^−4^). Unsupervised metacell-based methods performed poorly, with ArchR (0.223) and Cicero (0.247) achieving lowest mAUPRC values. Similar trends were observed in other low-sample cell types such as CD8 TEM1 (302 cells/8 metacells), CD8 TEM2 (363 cells/8 metacells), and CD16 Monocytes (521 cells/11 metacells), where BayesCNet again led in performance, followed by DIRECT-NET. Evaluation using AUROC produced consistent results (**Fig. 5b**). BayesCNet achieved the highest mAUROC values across all cell types, whereas Cicero, DIRECT-NET, and SnapATAC showed moderate performance, and ArchR and SCARlink approached random. BayesCNet, DIRECT-NET, and SnapATAC also yielded the highest number of evaluable marker genes (i.e., genes with ≥5 PCHi-C–overlapping predicted pairs) (**Supplementary Fig.5a**).

We next applied the methods to the HEMA scMultimodal dataset, which comprise eight hematopoietic progenitor populations, including CMP, GMP, HSC, LMPP, MPP, MEP, CLP and Mono. Progenitor cell types were matched to terminal PCHi-C profiles via the myeloid and lymphoid lineages (**Supplementary Table S5**), as described in *Methods*. For metacell-based methods, we used K=10 for KNN graph-based aggregation. BayesCNet again achieved the strongest and most consistent performance, ranking first across all eight cell types (**Fig. 5c**). The advantage was most evident in rare populations: BayesCNet attained mAUPRCs of 0.502 for Mono (64 cells/8 metacells) and 0.530 for CLP (78 cells/8 metacells), compared to ArchR (0.436) and SnapATAC (0.464) as respective second-best performers. These improvements were statistically significant (Wilcoxon p = 2.12×10^−11^ for CLP vs. SnapATAC; p = 1.13×10^−3^ for Mono vs. ArchR). SCARlink failed to detect any PCHi-C-supported interactions in these rare cell types. Even in more abundant populations such as CMP (502 cells/47 metacells) and GMP (402 cells/37 metacells), BayesCNet achieved top mAUPRCs (0.476 and 0.477), outperforming second-best performer DIRECT-NET (0.375 and 0.363) respectively. Notably, ArchR identified no PCHi-C-confirmed interactions in these cases. AUROC-based evaluation confirmed the same ranking (**Fig. 5d**), with BayesCNet leading in all cell types, followed by DIRECT-NET. Similarly, BayesCNet, DIRECT-NET, and SnapATAC produced the largest number of evaluable marker genes (i.e., genes with ≥5 predicted pairs overlapping PCHi-C loops) (**Supplementary Fig 5b**).

### Enhancer-gene linkages are enriched for trait heritability and GWAS variants

To assess the biological relevance of predicted EG linkages, we benchmarked BayesCNet and competing methods using Stratified linkage disequilibrium Score Regression (S-LDSC)^21^. S-LDSC partitions trait heritability across functional genomic annotations by integrating GWAS summary statistics with LD patterns. In this study, functional annotations corresponded to the top-quantile gene-linked peaks identified by each method. The goal was to evaluate which method most effectively identifies enhancer regions enriched for trait-associated heritability.

For the PBMC scMultiome dataset, we analyzed two systemic lupus erythematosus (SLE) GWAS from the NHGRI-EBI GWAS Catalog^22^ (GCST005831^23^ and GCST011097^24^). SLE was chosen because PBMCs are widely used to assess cellular composition and cell type-specific transcriptomic signatures in SLE ^25,26^. Gene-linked peaks were analyzed across nine immune cell types. S-LDSC analyses were conducted using the baselineLD v2.2 model, which incorporates 97 functional annotations to control for known genomic features (*Methods*). For GCST011097, BayesCNet achieved the highest enrichment in seven of nine cell types and reached statistical significance (p < 0.05) in all cell types (**Fig. 6a**). High enrichments were observed in adaptive immune populations, exceeding 100-fold in several cases, such as Naïve B (162×), CD4 Naïve (143×), CD8 Naïve (112×), CD4 TCM (110×), CD4 TEM (82×), and CD8 TEM1 (74×). Notably, Naïve B is the rarest population in the dataset, highlighting BayesCNet’s strength in low-sample regimes. BayesCNet also led in innate immune populations, including CD16 Mono (74.5×) and CD14 Mono (59.1×). BayesCNet’s performance is comparable to DIRECT-NET in CD8 TEM2 (101.2× vs. 111.5×) and to ArchR in CD14 Mono (59.1× vs. 64.6×). In contrast, SCARlink showed negligible enrichment across all cell types. SnapATAC exhibited intermediate performance but failed completely in CD4 Naïve. DIRECT-NET achieved high enrichment in CD8 TEM2 yet failed in Naïve B. In contrast, Cicero performed well in Naïve B (65.8×) but poorly in CD4 Naïve. ArchR yielded moderate enrichment in monocytes (CD14 Mono, 64.7×; CD16 Mono, 63.4×) but was weak elsewhere. Results were robust across multiple thresholds of gene-linked peaks (top 10%, 25%, and 50%) (**Supplementary Fig.6a**) and consistent with the other SLE GWAS study-GCST005831 (**Supplementary Fig. 6b**).

**Fig. 6.**
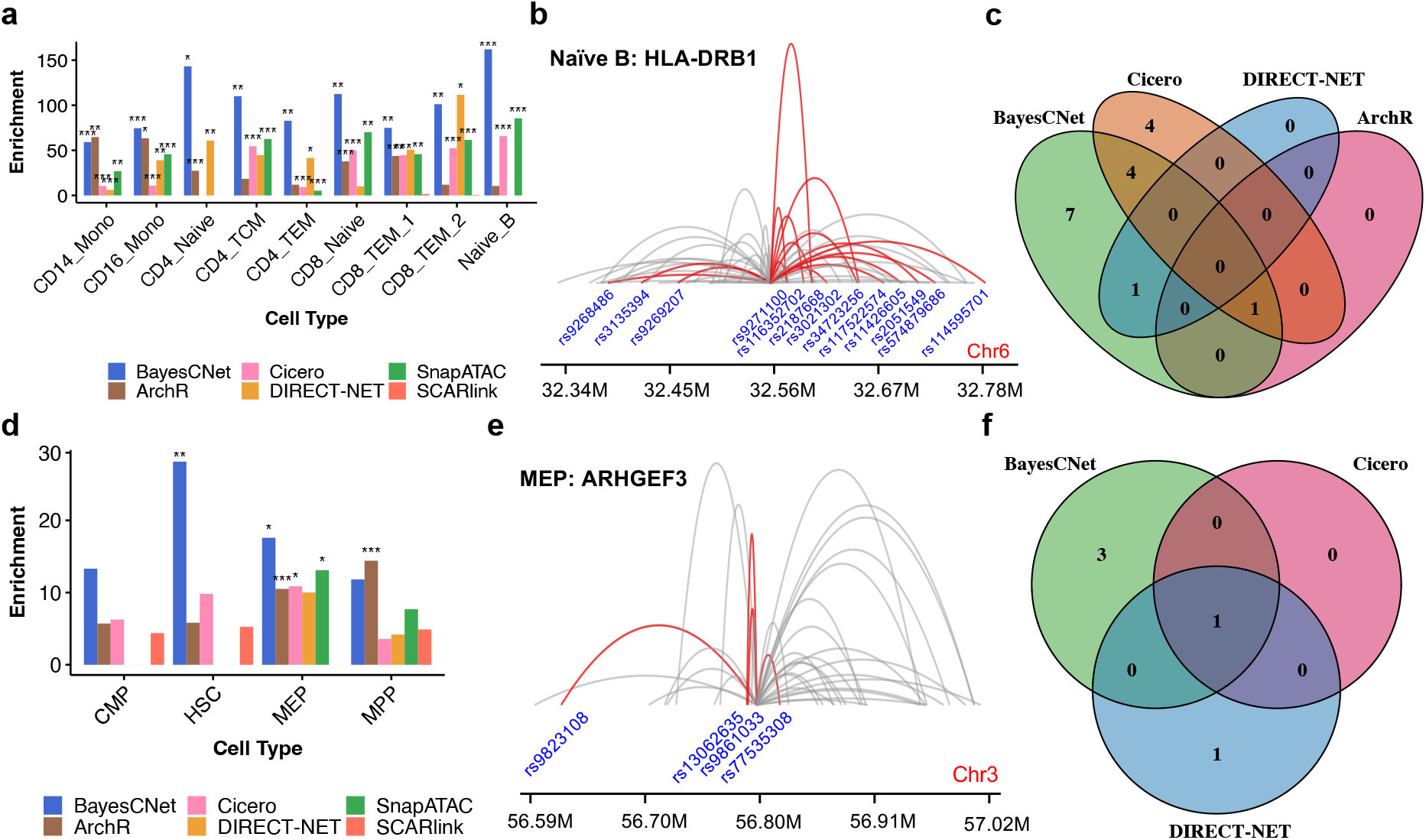
Evaluating biological relevance of cell type-specific enhancer-gene linkages using stratified linkage disequilibrium score regression (S-LDSC). **a**. S-LDSC enrichment for systemic lupus erythematosus (GWAS Catalog ID: GCST011097) based on enhancers from the top 25% of predicted enhancer–gene linkages identified by each method across cell types in PBMC scMultiome dataset. **b**. BayesCNet-identified Naïve B-specific enhancer–gene linkages colocalize with 13 SLE-associated SNPs at *HLA-DRB1* locus. All arcs denote BayesCNet-identified EG linkages. EG linkages overlapped with SLE-associated SNPs are highlighted in red. **c**. Venn diagram comparing SLE-associated SNPs in enhancers from EG linkages identified by all methods at *HLA-DRB1* locus **d**. S-LDSC enrichment for platelet volume (GWAS Catalog ID: GCST90028996) using the enhancers from top 25% of predicted enhancer–gene linkages from the HEMA scMultimodal dataset. **e**. BayesCNet-identified MEP-specific enhancer–gene linkages colocalize with 4 platelet volume-associated SNPs near *ARHGEF3*. **f**. Venn diagram comparing platelet volume-associated SNPs in these enhancers across methods at the *ARHGEF3* locus. Asterisks indicate statistical significance from S-LDSC analysis (p < 0.05; p < 0.01; p < 0.001; p < 0.0001).

We also evaluated the enrichment of significant SLE-associated GWAS SNPs (p < 5 × 10^−8^), obtained from NHGRI-EBI GWAS Catalog^22^, within EG linkages at *HLA-DRB1 and HLA-DQB1* locus, which is a well-established SLE risk region ^27,28^ (**Fig. 6b**). To ensure comparability, all methods were restricted to a ±250 kb window centered on the *HLA-DRB1* TSS when inferring EG linkages in Naïve B cells, the rarest PBMC population. BayesCNet identified a dense cluster of EG linkages overlapping 13 significant SLE-associated SNPs, outperforming Cicero (9 SNPs), DIRECT-NET (1 SNP), ArchR (1 SNP), and SnapATAC (0 SNPs). Of these, seven SNPs were uniquely detected by BayesCNet, while six were shared with other methods (**Fig. 6c**). Specifically, five SNPs overlapped between BayesCNet and Cicero, and the single SNPs detected by DIRECT-NET or ArchR were also identified by BayesCNet. Similarly, at the *HLA-DQB1* locus, BayesCNet again demonstrated stronger colocalization with significant SLE-associated SNPs than competing methods (**Supplementary Fig.7**). This locus-specific analysis highlights BayesCNet’s ability to interpret noncoding variants by linking them to downstream gene expression through EG linkages, thereby uncovering trait-associated regulatory activity at disease loci.

For the HEMA scMultimodal dataset, we analyzed platelet-related GWAS (platelet volume: GCST90028996^29^ and platelet count: GCST90468095^30^) from NHGRI-EBI GWAS Catalog^22^. Platelet traits were chosen because they directly reflect hematopoiesis, with platelets produced from megakaryocytes that differentiate from HSCs. Accordingly, we assessed four relevant progenitor types, including HSC, MPP, CMP and MEP (**Fig. 1b**). For platelet volume trait, BayesCNet achieved the highest enrichment in three of four populations (HSC, CMP, and MEP) and reached statistical significance in HSC and MEP (**Fig. 6d**). ArchR led in MPP but showed weaker enrichment elsewhere. Cicero yielded moderate enrichment, reaching significance only in MEP. SCARlink showed moderate enrichment in CMP, HSC and MPP but none in MEP. DIRECT-NET and SnapATAC performed moderately in MPP and MEP but failed in HSC or CMP. Overall, these results highlight BayesCNet’s superior ability to detect trait-relevant regulatory elements, while competing methods showed weaker or inconsistent enrichment. Importantly, enrichment patterns remained consistent across multiple percentile thresholds of gene-linked peaks (top 10%, 25%, and 50%), confirming the robustness of BayesCNet (**Supplementary Fig.8a**). Findings were further supported by a platelet count GWAS (GCST90468095) (**Supplementary Fig. 8b**).

Similar to the colocalization analysis on the SLE risk locus, we further evaluated colocalization of platelet-associated significant GWAS SNPs (p < 5 × 10^−8^) with inferred EG linkages from different methods in the HEMA scMultimodal dataset, focusing on MEPs, the rarest population and the direct precursors of platelet-producing megakaryocytes^31,32^. We also restricted all methods to a ±250 kb window centered on the *ARHGEF3* TSS, a gene with well-established roles in hematopoiesis and megakaryopoiesis^33,34^. BayesCNet identified four platelet count–associated SNPs, including three uniquely detected and one shared with Cicero and DIRECT-NET (**Fig. 6e, f**). In contrast, SnapATAC, ArchR, and SCARlink detected none. These findings demonstrate BayesCNet’s ability to uncover trait-associated regulatory interactions in rare progenitor populations, linking noncoding variants to their putative target genes. The colocalization analyses for both single cell datasets highlight BayesCNet’s broad utility in interpreting GWAS findings across diverse contexts, from immune loci such as *HLA-DRB1* associated with SLE to hematopoietic loci such as *ARHGEF3* associated with platelet traits.

### BayesCNet constructs cell type–specific TF–gene regulatory networks

We extended BayesCNet to construct cell type–specific TF–gene regulatory networks (TF-GRNs) by integrating three components: TF binding potential at gene-linked enhancers, predicted enhancer–gene linkages, and cell type–specific TF expression (*Methods*). TFs with high degree centrality are highlighted as potential key regulators. We first applied this framework to three PBMC cell types representing different abundance levels: Naïve B cells (low), CD4 TCM (medium), and CD14 monocytes (high). The top 20 TFs, ranked by degree centrality, in each network, varied substantially across cell types (**Fig. 7a**). Most TFs were cell type–specific (9 unique to Naïve B, 11 to CD4 TCM, and 10 to CD14 monocytes), while four (VEZF1, NFKB1, NFKB2, IRF1) were shared across all three (**Fig. 7b**). Notably, many of these top-ranked TFs correspond to well-established regulators. Three shared TFs, NFKB1, NFKB2 and IRF1, play central roles in immune regulation^35,36^. Naïve B-specific TFs, such as EBF1, SPIB, and POU2F2 (Oct2), are critical for B cell differentiation, survival and maintenance^37-40^. CD4 TCM-specific TFs, including NFATC2, KLF2, and MAF, have been implicated in T cell development and function^41-45^. In CD14 Mono, the top-ranked TF CEBPB (CCAAT/enhancer-binding protein), regulates myeloid cell differentiation and activates the CD14 promoter ^46^.

**Fig. 7.**
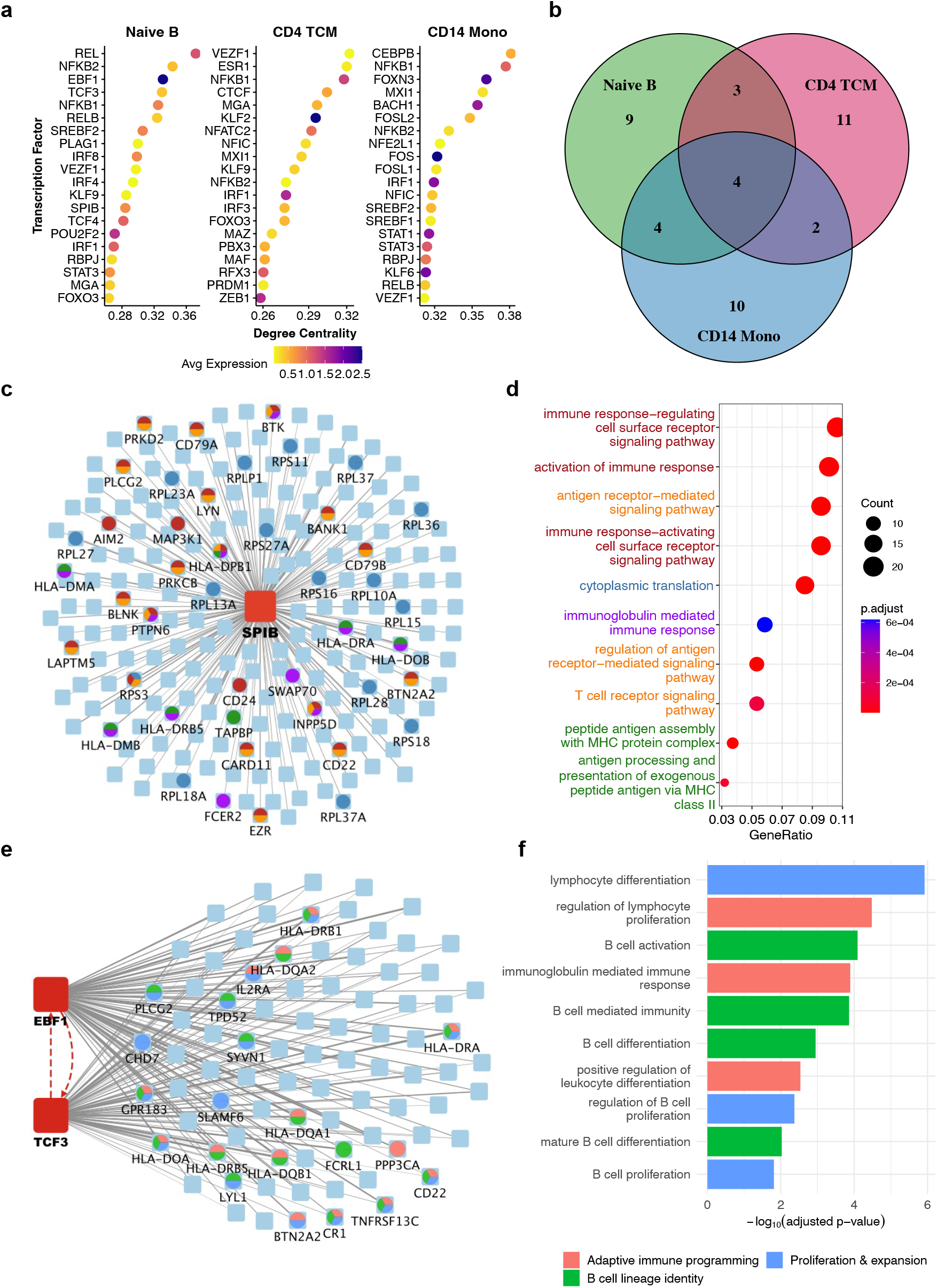
Cell type–specific TF–gene regulatory network (GRN) analysis in PBMC scMutiome dataset using BayesCNet. **a**. Top 20 transcription factors (TFs) ranked by degree centrality in Naive B cells, CD4 TCM, and CD14 monocytes. Node color indicates average TF expression. **b**. Venn diagram comparing top-ranked 20 TFs across the three cell types **c**. SPIB-hub TF-GRN in Naïve B cells. **d**, Gene Ontology (GO) enrichment analysis of SPIB target genes **e**. Two interacted TF-GRN between EBF1 and TCF3 TF hubs in Naive B cells. **f**. GO enrichment of EBF1–TCF3 shared gene targets.

As a representative case, we examined TF-GRN of top-ranked TFs in Naïve B. SPIB emerged as a high-centrality hub with extensive regulatory connections (**Fig. 7c**). A member of the ETS transcription factor family, SPIB is essential for immature B cell differentiation, functions as a DNA-binding TF, and regulates multiple components of the B-cell receptor signaling pathway^39,47^. Gene Ontology (GO) analysis of SPIB target genes revealed strong enrichment in immune response pathways, including immune response-regulating cell surface receptor signaling, activation of immune response, and antigen receptor-mediated signaling (**Fig. 7d**). These results underscore SPIB’s central role in orchestrating B cell-specific programs, likely in concert with its downstream target genes. Beyond single TF hubs, TF-GRN analysis also uncover regulatory interactions between multiple TF networks through TF–TF linkages. In Naïve B cells, EBF1 (Early B-cell Factor 1) and TCF3 (E2A) formed mutually connected TF hubs, where they are linked to each other, and their GRNs shared many target genes (**Fig. 7e**). This mutual regulatory relationship aligns with prior studies showing their coordinated function. For example, TCF3 activates EBF1^*48*^ and together they synergistically upregulate transcription of endogenous B cell–specific genes^49^, thereby playing important roles in early differentiation of B lymphocytes. Both factors also participate in a global pro-B cells regulatory network, binding regulatory elements at the Foxo1 locus and cooperating with Foxo1 interacted with putative enhancers in the Pax5 locus^50^. GO analysis of their shared target genes revealed strong enrichment for B-cell activation, differentiation, and proliferation (**Fig. 7f**). These findings further support the cooperative roles of EBF1 and E2A in regulating B lymphocyte differentiation.

We next applied BayesCNet to construct TF-GRNs in the HEMA scMultimodal dataset, focusing on three representative cell types, including Monocytes, GMPs and CLPs, to capture both rare and intermediate cell populations (**Supplementary Fig. 9a**). Comparative analysis of the top 20 TFs ranked by degree centrality in each network revealed both shared and cell type– specific regulators. Five TFs were common to all three lineages, while Monocytes and CLPs shared nine TFs, Monocytes and GMPs shared eight, and CLPs and GMPs shared eight. Each cell type also displayed distinct regulators, with GMPs harboring nine unique TFs and Monocytes and CLPs each having eight (**Supplementary Fig. 9b**). As a detailed case study, we examined REL (c-Rel), which emerged as a central hub in the monocyte-specific TF-GRN (**Supplementary Fig. 9c**). REL, a member of the NF-κB family, plays a central role in inflammasome activation, cytokine release, and cell survival in monocytes and macrophages^51^. It also acts as a transcriptional repressor in the maintenance of inflammatory homeostasis^52^ and has been implicated in regulating monocyte signature genes^53^. GO analysis of REL target genes confirmed significant enrichment for biological processes central to myeloid biology, including regulation of hematopoiesis, myeloid cell differentiation, and leukocyte differentiation (**Supplementary Fig. 9d**). These findings underscore BayesCNet’s ability to reveal biologically meaningful TF modules and highlight the importance of NF-κB/Rel signaling in shaping monocyte-specific regulatory programs. TF-GRN analysis also unveiled a TF-TF interaction module that links IRF1 and REL TF hubs (**Supplementary Fig. 9e**). IRF1 directly regulates REL and both TFs converged on a shared set of target genes. IRF1 is a regulator of class I and II MHC gene expression^54^ and orchestrates interferon-stimulated gene responses in human monocytes and macrophages^55^. Moreover, the activation of the NF-κB/IRF1 axis programs dendritic cells to promote antitumor immunity^56^. GO analysis of their shared target genes highlighted enrichment in antigen processing and presentation of peptide antigen via MHC class II, MHC class II protein complex assembly, and immunoglobulin-mediated immune responses (**Supplementary Fig. 9f**). Together, these results point to a cooperative regulatory circuit between IRF1 and REL in monocytes, linking interferon signaling with NF-κB–mediated immune activation.

## Discussion

We introduced BayesCNet, a Bayesian hierarchical framework, for mapping cell-type-specific EG linkages from single-cell data, applicable to paired RNA and ATAC, unpaired modalities, or ATAC alone. BayesCNet addresses a critical challenge in regulatory inference for rare cell types by borrowing information across related populations organized in a cell type hierarchy. By clustering cells into metacells via a KNN graph, BayesCNet mitigates sparsity and enables stable statistical inference of regulatory programs. EG linkages are inferred within a Bayesian regression framework that jointly models gene expression across all cell types as the response variables and accessibility of nearby ATAC peaks as the covariates, with structured priors capturing both cell type relationships (via cell hierarchies) and peak dependencies (via genomic proximity or co-accessibility). These priors can be derived from external biological knowledge or estimated directly from single-cell data. BayesCNet further integrates EG linkages with TF binding potential and cell type–specific TF expression to construct TF-GRNs, highlighting both shared and cell type–specific regulators of differentiation, activation, and disease processes.

Through comprehensive simulations, BayesCNet consistently outperformed state-of-the-art methods under diverse conditions, including different forms of cell type hierarchy (COT vs. CLT), varying sample sizes of rare cell types, and different topological positions of rare cell types within the hierarchy. Notably, BayesCNet showed the greatest advantage when rare populations were positioned at highly connected nodes, reflecting its ability to exploit cell type hierarchies. In real data applications, BayesCNet-inferred EG linkages were more concordant with cell type–specific PCHi-C interactions than other methods, in both PBMC scMutiome and HEMA scMutimodal datasets, which are two canonical systems representing ontology- and lineage-based hierarchies. S-LDSC further demonstrated that BayesCNet-identified enhancers were most enriched for trait-associated heritability, and BayesCNet uniquely captured disease-relevant GWAS SNPs within EG linkages for key immune and hematopoietic genes. Application to TF-GRN construction revealed both common and cell type–specific TFs, with enriched pathways consistent with established regulatory roles, underscoring BayesCNet’s biological interpretability.

There are several possible extensions for BayesCNet. In its current form, BayesCNet performed metacell aggregation of gene expression or chromatin accessibility to reduce sparsity in data modeling. Future implementations could incorporate zero-inflated distributions on the data modality directly to better capture the discrete and sparse nature of single-cell counts, thereby reformulating the framework into a Bayesian zero-inflated regression model. Although the present study focused on cells from a single biosample, BayesCNet can be generalized to integrate data across multiple biosamples, cohorts, or developmental stages. Such extensions could be achieved either through a two-step harmonization strategy prior to modeling or by explicitly incorporating biosample-level variation as covariates in the regression framework. In addition, BayesCNet could be extended to multi-condition settings by modeling condition-specific effects, thereby enabling inference of differential EG linkages across conditions. Given its demonstrated power, particularly in rare cell populations, BayesCNet will represent a broadly applicable tool for mapping cell type–specific regulatory programs and connecting GWAS variants to disease-relevant biology.

## Methods

### Data processing

We obtained the PBMC scMutiome data with co-assayed RNA and ATAC from 10x Genomics website and performed data processing by following the weighted nearest neighbor (WNN) analysis workflow in the Seurat vignette^57^. Briefly, gene expression and chromatin accessibility matrices were extracted and integrated into a multimodal Seurat object with ATAC features added using Signac^3^. Standard quality control filters were applied, followed by SCTransform^3^ normalization for RNA and TF–IDF/LSI dimensionality reduction for ATAC. A WNN graph was then constructed by combining RNA principal components and ATAC LSI embeddings, and cell clustering was performed. A summary of cell types and sample sizes is provided in **Supplementary Table S1**. For human hematopoietic differentiation (HEMA) scMutimodal data, which contains unpaired scRNA-seq and scATAC-seq of human hematopoietic cells^58^, we implemented an unsupervised matching strategy to align cells across modalities. Specifically, gene activity scores were computed from the ATAC modality using Signac, aggregating peak signals within a 2 kb window upstream of each gene’s TSS to generate a gene activity matrix *E*_*ATAC*_. A corresponding gene expression matrix *E*_*RNA*_ was obtained from RNA modality. Highly variable genes shared between modalities were retained, and the filtered matrices *E*_*ATAC*_ and *E*_*RNA*_ were concatenated along the cell dimension. The combined matrix was projected into a shared low-dimensional space using PCA. Pearson correlations were then calculated between ATAC-derived gene activity and RNA expression profiles using top 30 PCs. For each ATAC cell, the most correlated RNA cell was selected as its best match, yielding a one-to-one matched cell pair. The RNA expression and ATAC profiles of each matched pair were subsequently combined and treated as co-assayed in the same cell, generating what a pseudo scMultiome dataset. The distribution of aligned cell pairs across all cell types is shown in **Supplementary Table S2**.

## Model formulation

BayesCNet is a gene-centric model designed to infer gene-enhancer linkage by relating a gene’s promoter to nearby ATAC peaks, which represent candidate enhancers. Let *C* denote the number of cell types, *P* the unified set of ATAC peaks across all cell types, *N*_*c*_ the number of metacells in cell type *c*, and 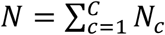 total number of metacells. For a given gene in cell type *c*, gene expression is modeled as a linear function of chromatin accessibility: *Y*_*c*_ = *X*_*c*_*B*_*c*_ + *E*_*c*_, where 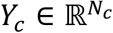 is the gene expression for cell type *c*, 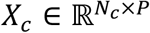 the ATAC profile for cell type *c*, and *B*_*c*_ = [*β*_*c*1,_*β*_*c*2_, …, *β*_*cP*_]^T^ ∈ ℝ^*P*^ the regression coefficients that quantify EG linkages. A large positive *β*_*ck*_ indicates a strong EG linkage between the gene and *k*-th ATAC peak in cell type *c*. To leverage cell type hierarchy and improve inference, especially for rare cell types, BayesCNet jointly models all *C* cell types using a block-diagonal regression framework: *Y* = *XB* + *E, E*∼*N*(0, σ^2^II_N_). Here, 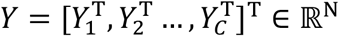 is the concatenated gene expression across all cell types. 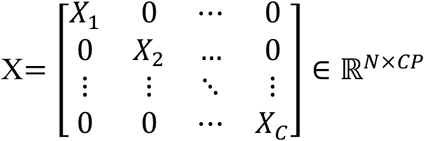is the block-diagonal design matrix of ATAC profiles across all cell types and 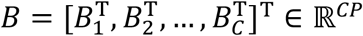 stacks the regression coefficients across peaks and cell types.

### Prior distribution on regression coefficients

To enable information sharing across both peaks and cell types, we reshape the coefficient vector *B* ∈ ℝ^*CP*^ into a matrix *B* ∈ ℝ^*P*×*C*^,where rows correspond to peaks and columns to cell types. We then place a matrix normal prior, *B*∼ℳ*N*_*P*×*C*_(0, Σ_1_, Σ_2_), which is equivalent to vec(*B*)∼*N*(0, Σ_2_ ⊗ Σ_1_). Here, Σ_1_ ∈ ℝ^*P*×*P*^ captures the peak dependencies, while Σ_2_ ∈ ℝ^*C*×*C*^ encodes cell type relationships. For conjugate inference, we introduce an auxiliary variable δ ∈ ℝ^*P*×*C*^ with δ∼ℳ*N*_*P*×*C*_(*δ*_0_, Σ_1_, II_*C*_), which yields the transformation 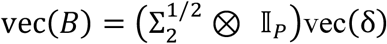. This reformulation decouples peak dependences (Σ_1_) from cell type relationships (Σ_2_), enabling efficient conjugate inference through the reduced model: 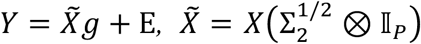, where *g* = vec(*δ*).

### Posterior inference

We employ conjugate Normal–Inverse-Gamma priors, which yield closed-form posterior updates. Specifically, we assume

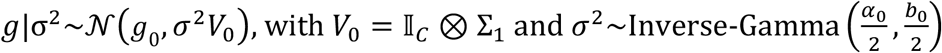

Posterior distribution of *g* is

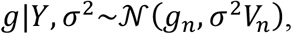

where 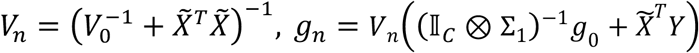 and 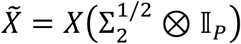

Posterior distribution of σ^2^ follows

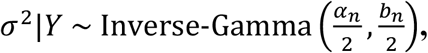

with 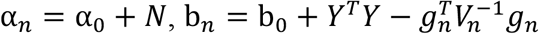

The posterior distribution of the regression coefficients *B* follows by transformation,

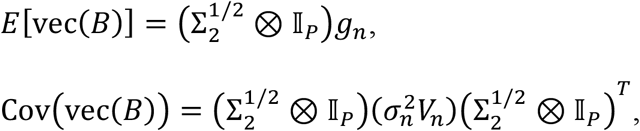

where 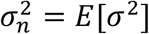. Reshaping the posterior mean yields *B* ∈ ℝ^*P*×*C*^, representing peak-level regression coefficient for each cell type. The posterior covariance decomposes into *C* block-diagonal submatrices Σ_*c*_ ∈ ℝ^*P*×*P*^, each representing peak–peak covariance within a cell type.

To quantify the strength of EG linkages, we define a Bayesian importance score *L*_*c*_ ∈ ℝ^*P*^ as

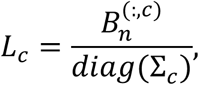

where 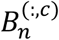 is the posterior mean and *diag*(Σ_*c*_) is the posterior variance of regression coefficients in cell type *c*. Larger positive entries in *L*_*c*_ indicate stronger evidence for EG linkages of corresponding peaks in cell type *c*. We set hyperparamers *g*_0_ = 0 to reflect no prior knowledge of peak effects and α_0_ = 1 and b_0_ = 1 to specify a weakly informative prior on the variance. More details of statistical inference can be found at Supplementary Notes.

### Estimation of Σ_1_ and Σ_2_

For each gene, we estimate Σ_1_ ∈ ℝ^*P* × *P*^, which captures the relationship among the *P* peaks, using an empirical Bayes approach motivated by Zellner’s g-prior^59^. Let *Z* ∈ ℝ^*N*×*P*^ be the vertically stacked design matrix across all cell types. We estimate 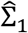 as 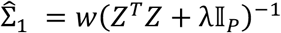, where λ is a small ridge penalty to ensure numerical stability (λ = 10^−6^ by default), and *w* is a data-driven scaling factor determined via Stein’s unbiased risk estimate (SURE). To characterize the cell type relationships, we obtain Σ_2_ ∈ ℝ^*C*×*C*^ either from prior biological knowledge or empirical estimation. When biological prior knowledge is available (e.g., COT and CLT), we first construct an adjacency matrix *A* ∈ ℝ^*C*×*C*^ by applying an exponential decay to the shortest path distances between cell types as *A*_*ij*_ = exp(−α*d*_*ij*_*), A*_*ii*_ = 1, where *d*_*ij*_ is the shortest path distance between cell types *i* and *j*, and α > 0, default 0.5, controls the decay rate with distance. Larger α induce more local similarity (i.e., only nearby cell types remain correlated), whereas smaller α allow more global similarity across distant cell types. From *A*, we compute the graph Laplacian *L* = *D* − *A*, where D is the diagonal degree matrix with entries *D*_*ii*_ = ∑_*j*_ *A*_*ij*_. For numerical stability, we regularize and invert the Laplacian as 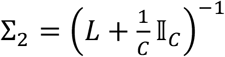,and then normalize Σ_2_ as 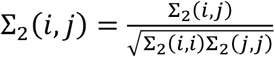 such that each diagonal entry equals 1. When biological prior knowledge is not available, we estimate Σ_2_ directly from the single-cell data. For each cell type, we extract the top 10 principal components from either RNA, ATAC, or a joint WNN embedding for each cell type. Pairwise distances between cell types are then computed using optimal transport with a Euclidean cost^60^,yielding a distance matrix *d*_0*T*_. We then transform these distances into a covariance using an exponential kernel: 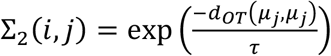, where *d*_0*T*_(*µ*_*j*_, *µ*_*j*_) is the OT distance between embeddings of the cell type *i* and *j*, and τ is a bandwidth parameter.

### Simulation studies

To benchmark BayesCNet against competing methods for inferring EG linkages, we conducted comprehensive simulation studies designed to evaluate (i) different forms of cell type hierarchy (COT vs. CLT), (ii) variations in tree topology, including placement of rare cell types, and (iii) sample size variations in the rare populations. Simulations were designed to reflect real single-cell data characteristics, incorporating (i) cell type hierarchies mimic ontology or lineage trees, (ii) peak–peak correlations, and (iii) empirical distributions from ATAC and RNA modalities. We further assessed the robustness of BayesCNet under priors derived from biological knowledge versus empirical estimation.

### Cell type hierarchy

Two types of hierarchies were modeled. COT, mimicking PBMCs and CLT, resembling hematopoietic differentiation. Both were encoded into adjacency matrices, followed by an exponential decay function to the shortest-path distances between cell types, which were then transformed into a covariance matrix Σ_2_. For peak-peak correlation, we selected 60 peaks and constructed Σ_1_. Approximately 10% of peaks pairs were randomly assigned correlations (*ρ*_*pos*_ = 0.5 or *ρ*_*neg*_ = −0.5), with all other pairs set to zero. Positive definiteness was enforced via eigenvalue adjustment.

### Generating ATAC profile and gene expression

we first simulated latent variable *Z* ∈ ℝ^*N*×*P*^from a multivariate normal distribution, vec(*Z*)∼*N*(0, Σ_2_ ⊗ Σ_1_), where Σ_1_ and Σ_2_ were estimated from real scATAC-seq (*Estimation of Σ* _1_*and Σ* _2_). This generated a large latent matrix 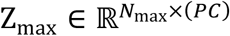, where *N*_max_ is the maximum number of cells per cell type. Cell type-specific latent matrices were then extracted as *Z*_*c*_ = *Z*_max_[1: *N*_*c*_, (*P*(*c* − 1) + 1): (*Pc*)]. Latent Gaussian variable *Z*_*c*_ were transformed to realistic ATAC counts *X*_*c*_ using Gaussian copula. Gaussian copula was chosen for its simplicity and proven ability to preserve correlation structure, which has been successfully applied to generate microbiome and single-cell RNA-seq count data^61,62^. Specifically, *Z*_*c*_ was mapped to uniforms via the normal cumulative distribution function (CDF) as *U*_*c*_ = Φ(*Z*_*c*_), which were then transformed into ATAC counts from a negative binomial distribution, using the inverse CDF as 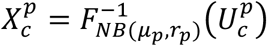, where μ_*p*_ and *r*_*p*_ are mean and dispersion parameter for peak *p*. Both parameters were empirically estimated from ATAC modality of PBMC scMultiome dataset by fitting negative binomial models to 5,000 randomly selected peaks, and simulation parameters were sampled from their empirical interquartile ranges.

We further converted the simulated single-cell ATAC cell counts into metacells. After filtering non-informative peaks, PCA was applied on *X*_*c*_ to obtain top 10 PCs, followed by K-means clustering to generate 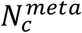 cell groups. The ATAC counts were summed within each cluster to generate a metacell-by-peak matrix 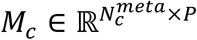. The full block-diagonal design matrix across all cell types was assembled as 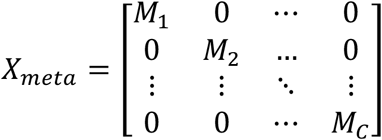. Regression coefficients *B* ∈ ℝ^*P*×*C*^ were sampled from a matrix normal distribution *B*∼ℳ*N*(0, Σ_1_, Σ_2_). Ground-truth EG linkages were defined as binary labels, with *B*_*jc*_ ≥ 0.5 indicating a true linkage between peak *j* and the given gene in cell type *c*. For single-cell methods (e.g. SnapATAC and SCARlink), which used single-cell ATAC and RNA counts as input, gene expression was simulated using a log-linear negative binomial model: μ = exp(*X*_c_vec(*B*)), *Y*_c_∼*NB*(μ, *θ*), where mean μ and dispersion *θ* were estimated from RNA modality of PBMC scMultiome dataset. For metacell-based methods (e.g., BayesCNet, Cicero, ArchR, DIRECT-NET), gene expression was simulated using a linear gaussian model: *Y*_meta_ = *X*_meta_vec(*B*) + *ε, ε*∼*N*(0, *I*).

### Prior specification for cell type hierarchy

In the simulation studies, we compared BayesCNet’s performance under two different prior specification of cell type relationship, encoded in Σ_2_, each representing a different level of prior knowledge. When biological prior was available (e.g., COT or CLT), we applied an exponential decay function to pairwise shortest-path distances among cell types, which is further transformed into Σ_2_. Without such prior, we instead estimated Σ_2_ empirically from the single-cell data. Specifically, we computed pairwise optimal transport distances between cell types using a Euclidean cost, based on embeddings from RNA, ATAC, or joint WNN space, and transformed these distances into into Σ_2_ via an exponential kernel. This dual approach enabled systematic evaluation of BayesCNet’s robustness under varying degrees of prior information.

### Inferring enhancer-gene linkages in real data application

We benchmarked BayesCNet against competing methods for inferring GE linkages using two real datasets: 10X PBMC single-cell multiome dataset, referred to “PBMC scMultiome dataset” and the human adult hematopoietic differentiation dataset, referred to “HEMA scMultimodal dataset”. Data processing steps for both datasets are detailed in the *Data processing*. The PBMC scMultiome dataset was chosen because it encompasses a broad set of well-characterized immune cell types that can be organized into a COT, representing a hierarchical classification of cell types. PBMC COT was adapted from previous studies^63,64^. This dataset also provides an illustrative case for benchmark when RNA and ATAC are co-assayed in the same cell. The HEMA scMultimodal dataset was selected because hematopoietic differentiation represents a canonical example of a lineage tree, tracing trajectories from hematopoietic stem cells (HSCs) through progenitors to terminally differentiated lineages. HEMA CLT was constructed based on established hematopoietic differentiation pathways^65,66^. This dataset also exemplifies a benchmark case when RNA and ATAC are profiled in different cells. For both datasets, the cell type hierarchies were converted into structured priors following the procedure described in the *Cell type hierarchy*. To ensure fair evaluation across methods, we adopted a gene selection procedure similar to DIRECT-NET, focusing on cell type-specific marker genes. Marker genes, expected to have distinct regulatory landscapes, were identified by differential expression analysis comparing each cell type against all others using Seurat’s *FindAllMarkers* function with parameters set to *only*.*pos=TRUE, min*.*pct=0*.*1* and *logfc*.*threshold=0*.*1*. The analysis was restricted to protein-coding genes only.

### Benchmarking methods and performance evaluation

We compared BayesCNet with five state-of-the-art methods that span supervised and unsupervised learning framework and operate at either the single-cell or metacell level. In terms of input resolution, BayesCNet, Cicero, ArchR, and DIRECT-NET use metacell-level aggregated data, whereas SCARlink and SnapATAC operate directly on single-cell data. Regarding learning paradigm, Cicero and ArchR are unsupervised approaches, while DIRECT-NET, SCARlink, SnapATAC, and BayesCNet are supervised. For real data analyses, all methods were implemented following their published pipelines with default parameters. Specifically, Cicero identifies co-accessible regions within a genomic window using scATAC-seq data. To reduce sparsity, it groups similar cells into metacells using a KNN graph and computes co-accessibility scores using a graphical LASSO-regularized correlation matrix, which links peaks to gene promoters. ArchR aggregates similar single cells into pseudo-bulk replicates and estimates peak co-accessibility by computing Pearson correlations of log2-normalized accessibility across these replicates within the defined genomic window. A high correlation indicates a strong EG linkage. DIRECT-NET models gene expression as a function of ATAC accessibility using an XGBoost model, with metacell-level aggregation using KNN graph. Feature importance scores from XGBoost represent confidence in EG linkages. SCARlink employs regularized Poisson regression to model gene expression as a function of ATAC accessibility, with coefficients constrained to be non-negative so that only positive regulatory associations are inferred.

SnapATAC applies logistic regression, regressing binary peak accessibility on gene expression for each gene–peak pair. For SCARlink and SnapATAC, a large positive coefficient indicates a strong EG linkage. For these approaches, EG linkages were inferred independently within each cell type. For BayesCNet, we follow the pipeline outlined in **Fig. 2**, considering only peaks with positive Bayesian importance scores as potential EG linkages. In our analyses, all approaches were applied to peaks or tiles in a window of ±250 kb windows around TSS. For simulation studies, all methods were applied using their core algorithm for comparison. For each method and cell type, we assessed predictive accuracy using the area under the precision–recall curve (AUPRC) and the area under the receiver operating characteristic curve (AUROC).

### Evaluation with Promoter Capture Hi-C data

For both PBMC multiome and HEMA multimodal datasets, we evaluated predicted cell type-specific EG linkages against a previously reported PCHi-C dataset^20^. This dataset includes high-confident chromatin loops (Chicago score ≥ 5) across multiple cell types (**Supplementary Table 3**). Because the evaluation is cell type-specific, we performed cell type matching between PCHi-C dataset and single-cell dataset. For the PBMC scMultiome dataset, cell types with exact correspondence in PCHi-C (e.g., CD4 Naïve) were directly matched. For PBMC cell types without exact counterparts, we mapped them to their broader parent cell types in PBMC COT and used those parent categories for matching to PCHi-C. For example, CD14 and CD16 monocytes in PBMC scMultiome dataset were both matched to the “monocyte” category in PCHi-C dataset. For the HEMA scMultimodal dataset, which primarily contains progenitors, we mapped each progenitor to terminal lineages in PCHi-C based on lineage identity. For example, CLP (common lymphoid progenitor) was matched to T cells and B cells, while CMP (common myeloid progenitor) was matched to monocytes, macrophages, neutrophils, megakaryocytes, and erythroid cells. Full mapping pairs are provided in **Supplementary Tables 4–5**. To ensure genome coordinate compatibility, ATAC-seq peaks (hg38) were converted to the hg19 reference using the UCSC liftOver file (hg38ToHg19.over.chain) and the *rtracklayer* R package. A predicted EG linkage was considered supported by PCHi-C if one end (i.e.gene promoter or peak) was within 1 kb of one loop anchor and the other end within 1 kb of the other anchor of a PCHi-C loop. Using this criterion, we assigned binary labels to predicted EG linkages. Model performance was evaluated by comparing binary labels against Bayesian importance score from BayesCNet or other scores from competing methods, using AUPRC and AUROC as metrics. For robustness, we restricted evaluation to marker genes with at least five predicted EG linkages overlapping PCHi-C loops. Although all methods began with the same set of marker genes for each cell type, their distinct predicted EG linkages led to separate filtering and method-specific evaluation gene sets.

### Stratified LD score regression (S-LDSC) analysis

We applied S-LDSC to quantify the heritability enrichment of cell type–specific regulatory elements across all methods. For each cell type, gene-linked peaks (or tiles) from the top-quantile EG linkages identified by each method were used as functional genomic annotations for S-LDSC analysis^21^. S-LDSC estimates the contribution of genomic annotations to the heritability of complex traits by modeling GWAS summary statistics as a function of LD scores, allowing partitioning of SNP-based heritability across functional categories. LD scores were then computed for these annotated SNPs using European reference genotypes from the 1000 Genomes Project Phase 3 panel^67^, capturing the local LD structure. GWAS test statistics were regressed on these LD scores in a joint model that included the focal enhancer annotation alongside 97 baseline annotations from the baselineLD v2.2 model. Analyses were restricted to autosomal chromosomes, HapMap3 SNPs outside the MHC region, and used precomputed regression weights to account for heteroskedasticity and LD-dependent variance. This procedure was repeated across all combinations of methods and cell types, and performance was evaluated based on heritability enrichment values and their statistical significance.

### Cell Type-Specific TF-Gene Network Construction

To construct cell type-specific TF-GRNs, we developed an integrative computational framework, which combines three key components: TF binding potential at peaks, gene–peak regulatory links predicted by BayesCNet and cell type-specific TF expression level. The GRN framework proceeds in three key steps.

**TF motif score** For each gene, we selected the gene-linked peaks from the top quantile of BayesCNet-predicted EG linkages. Motif scan was performed on these peaks (i.e. candidate enhancers) for TF motifs using *motifmatchr* R package with position weight matrices (PWMs) from JASPAR2020^68^. Each enhancer-TF pair yielded an observed motif score (S_obs_). To control for sequence bias, we generated an empirical background distribution separately for each peak by sampling 1,000 background peaks (n_bg_=1000) matched for GC content and length of the target peak using chromVAR^69^. Motif scanning was then performed on these background peaks to obtain a background scores S_bg_. For each enhancer-TF pair, an empirical p-value was computed as 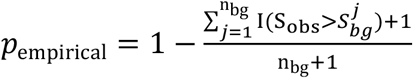. These p-values were transformed into normalized TF motif scores as *M* = −log_10_(*p*_empirical_).

**TF–enhancer–gene triplets** Using the normalized TF motif scores, we first identified significant TF-enhancer pairs (*t, e*) (*p*_empirical_<0.05). These pairs were then linked to BayesCNet-inferred EG linkages (*e, g*), forming TF–enhancer–gene triplets (*t, e, g*). For each triplet, we defined a triplet score as the geometric mean of three components as Score_*teg*_ = (*M*_*te*_ · *L*_*eg*_ · *E*_*t*_) ^1/3^, where *M*_*te*_ is the normalized TF motif score for enhancer *e, L*_*eg*_ is the Bayesian important score linking enhancer *e* to gene *g*, and *E*_*t*_ is the average expression of TF *t* in the corresponding cell type.

### TF-gene regulatory network

After computing triplet scores for all (*t, e, g*) combinations, we collapsed them to construct cell type–specific TF-GRNs. For each TF-gene pair (*t, g*), multiple enhancers may contribute. In such cases, the TF-gene edge weight was defined as the sum of the triplet scores across all enhancers associated with gene *g* as 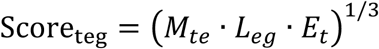, where *E*_*t*g_ denotes the set of enhancers linking TF *t* to gene *g*. The resulting weighted edges were assembled into TF-GRNs for each cell type and degree centrality was used to identify top-ranked TFs that act as potential key regulatory hubs.

## Data availability

The scRNA-seq and scATAC-seq data for human hematopoietic differentiation from Buenrostro et al. are provided from Zenodo https://zenodo.org/record/7879228. The single-cell multiome data for PBMC are available at 10X Genomics website. Promoter Capture Hi-C in 17 human primary blood cell types are provided from https://osf.io/u8tzp/

## Code availability

https://github.com/lichen-lab/BayesCNet

## Acknowledgement

This study was supported by National Institute of Health (NIH) grants R35GM142701 to L.C.

## References

1 Long, H. K., Prescott, S. L. & Wysocka, J. Ever-Changing Landscapes: Transcriptional Enhancers in Development and Evolution. Cell 167, 1170–1187 (2016). 10.1016/j.cell.2016.09.018

2 Sur, I. & Taipale, J. The role of enhancers in cancer. Nat Rev Cancer 16, 483–493 (2016). 10.1038/nrc.2016.62

3 Lenhard, B., Sandelin, A. & Carninci, P. Metazoan promoters: emerging characteristics and insights into transcriptional regulation. Nat Rev Genet 13, 233–245 (2012). 10.1038/nrg3163

4 Maurano, M. T. et al. Systematic localization of common disease-associated variation in regulatory DNA. Science 337, 1190–1195 (2012). 10.1126/science.1222794

5 Lieberman-Aiden, E. et al. Comprehensive mapping of long-range interactions reveals folding principles of the human genome. Science 326, 289–293 (2009). 10.1126/science.1181369

6 Fullwood, M. J. et al. An oestrogen-receptor-alpha-bound human chromatin interactome. Nature 462, 58–64 (2009). 10.1038/nature08497

7 Mumbach, M. R. et al. HiChIP: efficient and sensitive analysis of protein-directed genome architecture. Nat Methods 13, 919–922 (2016). 10.1038/nmeth.3999

8 Boix, C. A., James, B. T., Park, Y. P., Meuleman, W. & Kellis, M. Regulatory genomic circuitry of human disease loci by integrative epigenomics. Nature 590, 300–307 (2021). 10.1038/s41586-020-03145-z

9 Pliner, H. A. et al. Cicero Predicts cis-Regulatory DNA Interactions from Single-Cell Chromatin Accessibility Data. Molecular Cell 71, 858-871.e858 (2018). 10.1016/j.molcel.2018.06.044

10 Granja, J. M. et al. ArchR is a scalable software package for integrative single-cell chromatin accessibility analysis. Nature Genetics 53, 403–411 (2021). 10.1038/s41588-021-00790-6

11 Zhang, L., Zhang, J. & Nie, Q. DIRECT-NET: An efficient method to discover cis-regulatory elements and construct regulatory networks from single-cell multiomics data. Sci Adv 8, eabl7393 (2022). 10.1126/sciadv.abl7393

12 Fang, R. et al. Comprehensive analysis of single cell ATAC-seq data with SnapATAC. Nature Communications 12, 1337 (2021). 10.1038/s41467-021-21583-9

13 Mitra, S. et al. Single-cell multi-ome regression models identify functional and disease-associated enhancers and enable chromatin potential analysis. Nature Genetics 56, 627–636 (2024). 10.1038/s41588-024-01689-8

14 Hu, Z., Przytycki, P. F. & Pollard, K. S. CellWalker2: Multi-omic discovery using hierarchical cell type relationships. Cell Genom 5, 100886 (2025). 10.1016/j.xgen.2025.100886

15 Goodman, W. A. et al. KLF6 contributes to myeloid cell plasticity in the pathogenesis of intestinal inflammation. Mucosal Immunol 9, 1250–1262 (2016). 10.1038/mi.2016.1

16 Stuart, T. et al. Comprehensive Integration of Single-Cell Data. Cell 177, 1888–1902 e1821 (2019). 10.1016/j.cell.2019.05.031

17 Stuart, T., Srivastava, A., Madad, S., Lareau, C. A. & Satija, R. Single-cell chromatin state analysis with Signac. Nat Methods 18, 1333–1341 (2021). 10.1038/s41592-021-01282-5

18 Daugherty, A. C. et al. Chromatin accessibility dynamics reveal novel functional enhancers in C. elegans. Genome Res 27, 2096–2107 (2017). 10.1101/gr.226233.117

19 Johansen, N. J. et al. Evaluating methods for the prediction of cell-type-specific enhancers in the mammalian cortex. Cell Genom 5, 100879 (2025). 10.1016/j.xgen.2025.100879

20 Javierre, B. M. et al. Lineage-Specific Genome Architecture Links Enhancers and Non-coding Disease Variants to Target Gene Promoters. Cell 167, 1369-1384.e1319 (2016). 10.1016/j.cell.2016.09.037

21 Finucane, H. K. et al. Partitioning heritability by functional annotation using genomewide association summary statistics. Nat Genet 47, 1228–1235 (2015). 10.1038/ng.3404

22 Cerezo, M. et al. The NHGRI-EBI GWAS Catalog: standards for reusability, sustainability and diversity. Nucleic Acids Res 53, D998–D1005 (2025). 10.1093/nar/gkae1070

23 Julia, A. et al. Genome-wide association study meta-analysis identifies five new loci for systemic lupus erythematosus. Arthritis Res Ther 20, 100 (2018). 10.1186/s13075-018-1604-1

24 Wang, Y. F. et al. Identification of 38 novel loci for systemic lupus erythematosus and genetic heterogeneity between ancestral groups. Nat Commun 12, 772 (2021). 10.1038/s41467-021-21049-y

25 Herrada, A. A. et al. Innate Immune Cells’ Contribution to Systemic Lupus Erythematosus. Front Immunol 10, 772 (2019). 10.3389/fimmu.2019.00772

26 Perez, R. K. et al. Single-cell RNA-seq reveals cell type-specific molecular and genetic associations to lupus. Science 376, eabf1970 (2022). 10.1126/science.abf1970

27 Connolly, J. J. & Hakonarson, H. Role of cytokines in systemic lupus erythematosus: recent progress from GWAS and sequencing. J Biomed Biotechnol 2012, 798924 (2012). 10.1155/2012/798924

28 Grumet, F. C., Coukell, A., Bodmer, J. G., Bodmer, W. F. & McDevitt, H. O. Histocompatibility (HL-A) antigens associated with systemic lupus erythematosus. A possible genetic predisposition to disease. N Engl J Med 285, 193–196 (1971). 10.1056/NEJM197107222850403

29 Loh, P. R., Kichaev, G., Gazal, S., Schoech, A. P. & Price, A. L. Mixed-model association for biobank-scale datasets. Nat Genet 50, 906–908 (2018). 10.1038/s41588-018-0144-6

30 Loya, H., Kalantzis, G., Cooper, F. & Palamara, P. F. A scalable variational inference approach for increased mixed-model association power. Nat Genet 57, 461–468 (2025). 10.1038/s41588-024-02044-7

31 Deutsch, V. R. & Tomer, A. Megakaryocyte development and platelet production. Br J Haematol 134, 453–466 (2006). 10.1111/j.1365-2141.2006.06215.x

32 Psaila, B. et al. Single-cell profiling of human megakaryocyte-erythroid progenitors identifies distinct megakaryocyte and erythroid differentiation pathways. Genome Biol 17, 83 (2016). 10.1186/s13059-016-0939-7

33 Eicher, J. D. et al. Platelet-Related Variants Identified by Exomechip Meta-analysis in 157,293 Individuals. Am J Hum Genet 99, 40–55 (2016). 10.1016/j.ajhg.2016.05.005

34 Zou, S. et al. SNP in human ARHGEF3 promoter is associated with DNase hypersensitivity, transcript level and platelet function, and Arhgef3 KO mice have increased mean platelet volume. PLoS One 12, e0178095 (2017). 10.1371/journal.pone.0178095

35 Beinke, S. & Ley, S. C. Functions of NF-kappaB1 and NF-kappaB2 in immune cell biology. Biochem J 382, 393–409 (2004). 10.1042/BJ20040544

36 Feng, H., Zhang, Y. B., Gui, J. F., Lemon, S. M. & Yamane, D. Interferon regulatory factor 1 (IRF1) and anti-pathogen innate immune responses. PLoS Pathog 17, e1009220 (2021). 10.1371/journal.ppat.1009220

37 Hodson, D. J. et al. Regulation of normal B-cell differentiation and malignant B-cell survival by OCT2. Proc Natl Acad Sci U S A 113, E2039–2046 (2016). 10.1073/pnas.1600557113

38 Nechanitzky, R. et al. Transcription factor EBF1 is essential for the maintenance of B cell identity and prevention of alternative fates in committed cells. Nat Immunol 14, 867–875 (2013). 10.1038/ni.2641

39 Horiuchi, S. et al. SpiB regulates the expression of B-cell-related genes and increases the longevity of memory B cells. Front Immunol 14, 1250719 (2023). 10.3389/fimmu.2023.1250719

40 Emslie, D. et al. Oct2 enhances antibody-secreting cell differentiation through regulation of IL-5 receptor alpha chain expression on activated B cells. J Exp Med 205, 409–421 (2008). 10.1084/jem.20072049

41 Macian, F. NFAT proteins: key regulators of T-cell development and function. Nat Rev Immunol 5, 472–484 (2005). 10.1038/nri1632

42 Weinreich, M. A. et al. KLF2 transcription-factor deficiency in T cells results in unrestrained cytokine production and upregulation of bystander chemokine receptors. Immunity 31, 122–130 (2009). 10.1016/j.immuni.2009.05.011

43 Bopp, T. et al. NFATc2 and NFATc3 transcription factors play a crucial role in suppression of CD4+ T lymphocytes by CD4+ CD25+ regulatory T cells. J Exp Med 201, 181–187 (2005). 10.1084/jem.20041538

44 Gabrysova, L. et al. c-Maf controls immune responses by regulating disease-specific gene networks and repressing IL-2 in CD4(+) T cells. Nat Immunol 19, 497–507 (2018). 10.1038/s41590-018-0083-5

45 Dienz, O. et al. Accumulation of NFAT mediates IL-2 expression in memory, but not naive, CD4+ T cells. Proc Natl Acad Sci U S A 104, 7175–7180 (2007). 10.1073/pnas.0610442104

46 Pan, Z., Hetherington, C. J. & Zhang, D. E. CCAAT/enhancer-binding protein activates the CD14 promoter and mediates transforming growth factor beta signaling in monocyte development. J Biol Chem 274, 23242–23248 (1999). 10.1074/jbc.274.33.23242

47 Willis, S. N. et al. Environmental sensing by mature B cells is controlled by the transcription factors PU.1 and SpiB. Nat Commun 8, 1426 (2017). 10.1038/s41467-017-01605-1

48 Roessler, S. et al. Distinct promoters mediate the regulation of Ebf1 gene expression by interleukin-7 and Pax5. Mol Cell Biol 27, 579–594 (2007). 10.1128/MCB.01192-06

49 O’Riordan, M. & Grosschedl, R. Coordinate regulation of B cell differentiation by the transcription factors EBF and E2A. Immunity 11, 21–31 (1999). 10.1016/s1074-7613(00)80078-3

50 Lin, Y. C. et al. A global network of transcription factors, involving E2A, EBF1 and Foxo1, that orchestrates B cell fate. Nat Immunol 11, 635–643 (2010). 10.1038/ni.1891

51 Mussbacher, M., Derler, M., Basilio, J. & Schmid, J. A. NF-kappaB in monocytes and macrophages - an inflammatory master regulator in multitalented immune cells. Front Immunol 14, 1134661 (2023). 10.3389/fimmu.2023.1134661

52 de Jesus, T. J. & Ramakrishnan, P. NF-kappaB c-Rel Dictates the Inflammatory Threshold by Acting as a Transcriptional Repressor. iScience 23, 100876 (2020). 10.1016/j.isci.2020.100876

53 Li, T. et al. c-Rel-dependent monocytes are potent immune suppressor cells in cancer. J Leukoc Biol 112, 845–859 (2022). 10.1002/JLB.1MA0422-518RR

54 Hobart, M., Ramassar, V., Goes, N., Urmson, J. & Halloran, P. F. IFN regulatory factor-1 plays a central role in the regulation of the expression of class I and II MHC genes in vivo. J Immunol 158, 4260–4269 (1997).

55 Song, R. et al. IRF1 governs the differential interferon-stimulated gene responses in human monocytes and macrophages by regulating chromatin accessibility. Cell Rep 34, 108891 (2021). 10.1016/j.celrep.2021.108891

56 Ghislat, G. et al. NF-kappaB-dependent IRF1 activation programs cDC1 dendritic cells to drive antitumor immunity. Sci Immunol 6 (2021). 10.1126/sciimmunol.abg3570

57 Hao, Y. et al. Integrated analysis of multimodal single-cell data. Cell 184, 3573–3587 e3529 (2021). 10.1016/j.cell.2021.04.048

58 Buenrostro, J. D. et al. Integrated Single-Cell Analysis Maps the Continuous Regulatory Landscape of Human Hematopoietic Differentiation. Cell 173, 1535-1548.e1516 (2018). 10.1016/j.cell.2018.03.074

59 Zellner, A. in Bayesian Inference and Decision Techniques: Essays in Honor of Bruno de Finetti (ed P. Goel and A. Zellner) 233–243 (Elsevier Science, 1986).

60 Schiebinger, G. et al. Optimal-Transport Analysis of Single-Cell Gene Expression Identifies Developmental Trajectories in Reprogramming. Cell 176, 1517 (2019). 10.1016/j.cell.2019.02.026

61 Kurtz, Z. D. et al. Sparse and compositionally robust inference of microbial ecological networks. PLoS Comput Biol 11, e1004226 (2015). 10.1371/journal.pcbi.1004226

62 Sun, T., Song, D., Li, W. V. & Li, J. J. scDesign2: a transparent simulator that generates high-fidelity single-cell gene expression count data with gene correlations captured. Genome Biol 22, 163 (2021). 10.1186/s13059-021-02367-2

63 Jiang, Y., Chen, Z., Han, N., Shang, J. & Wu, A. sc-ImmuCC: hierarchical annotation for immune cell types in single-cell RNA-seq. Frontiers in Immunology 14, 1223471 (2023). 10.3389/fimmu.2023.1223471

64 Huang, L. et al. A statistical method for association analysis of cell type compositions. Statistics in biosciences 13, 373–385 (2021). 10.1007/s12561-020-09293-0

65 Corces, M. R. et al. Lineage-specific and single-cell chromatin accessibility charts human hematopoiesis and leukemia evolution. Nature Genetics 48, 1193–1203 (2016). 10.1038/ng.3646

66 Ulirsch, J. C. et al. Interrogation of human hematopoiesis at single-cell and single-variant resolution. Nature Genetics 51, 683–693 (2019). 10.1038/s41588-019-0362-6

67 Genomes Project, C. et al. A global reference for human genetic variation. Nature 526, 68–74 (2015). 10.1038/nature15393

68 Fornes, O. et al. JASPAR 2020: update of the open-access database of transcription factor binding profiles. Nucleic Acids Res 48, D87–D92 (2020). 10.1093/nar/gkz1001

69 Schep, A. N., Wu, B., Buenrostro, J. D. & Greenleaf, W. J. chromVAR: inferring transcription-factor-associated accessibility from single-cell epigenomic data. Nat Methods 14, 975–978 (2017). 10.1038/nmeth.4401

