## Supplementary file for "BayesCNet: Bayesian inference for cell type-specific regulatory networks leveraging cell type hierarchy in single-cell data"

September 11, 2025

### Supplementary Notes

BayesCNet is a gene-centric model designed to infer gene-enhancer linkage by relating a gene's promoter to nearby ATAC peaks, which represent candidate enhancers. Let $C$ denote the number of cell types and $P$ the number of peaks. Let $N = \sum_{c=1}^{C} N_c$ be the total number of metacells, where $N_c$ is the number of metacells in cell type $c$.

#### BayesCNet model formularization

Let $\mathbf{Y} \in \mathbb{R}^N$ be gene expression values across all metacells, stacked by cell type

$$\mathbf{Y} = \begin{bmatrix} Y_1 \\ Y_2 \\ \vdots \\ Y_C \end{bmatrix}, \quad Y_c \in \mathbb{R}^{N_c}$$

Let $\mathbf{X} \in \mathbb{R}^{N \times CP}$ be the block-diagonal design matrix, where each diagonal block $\mathbf{X}_c \in \mathbb{R}^{N_c \times P}$ contains the accessibility values for the $P$ peaks in cell type $c$

$$\mathbf{X} = \begin{bmatrix} \mathbf{X}_1 & 0 & \cdots & 0 \\ 0 & \mathbf{X}_2 & \cdots & 0 \\ \vdots & \vdots & \ddots & \vdots \\ 0 & 0 & \cdots & \mathbf{X}_C \end{bmatrix}$$

Let $\mathbf{B} \in \mathbb{R}^{CP}$ the regression coefficients, stacked by the cell type-specific regression coefficients $\boldsymbol{\beta}_c \in \mathbb{R}^P$

$$\mathbf{B} = \begin{bmatrix} \boldsymbol{\beta}_1 \\ \boldsymbol{\beta}_2 \\ \vdots \\ \boldsymbol{\beta}_C \end{bmatrix}, \quad \boldsymbol{\beta}_c \in \mathbb{R}^P$$

The joint regression model is given by

$$\mathbf{Y} = \mathbf{X}\mathbf{B} + \boldsymbol{E}, \quad \boldsymbol{E} \sim \mathcal{N}(0, \sigma^2 \mathbf{I}_N) \tag{1}$$

#### Structure prior on regression coefficients

To encourage peak dependencies and cell type relationships, we reshape $\mathbf{B}$ into $\mathbf{B}_{P \times C}$, with rows representing peaks and columns representing cell types, and place a matrix normal prior on $\mathbf{B}_{P \times C}$

$$\mathbf{B} \sim \mathcal{MN}_{P \times C}(\mathbf{B}_0, \Sigma_1, \Sigma_2)$$

where $\Sigma_1 \in \mathbb{R}^{P \times P}$ models peak dependencies and $\Sigma_2 \in \mathbb{R}^{C \times C}$ captures cell type relationships. Equivalently, a vector form of $\mathbf{B}$, $\text{vec}(\mathbf{B}) \in \mathbb{R}^{PC}$, follows a multivariate normal distribution (MNN)

$$\text{vec}(\mathbf{B}) \sim \mathcal{N}(\mathbf{b}_0, \Sigma_2 \otimes \Sigma_1)$$

To facilitate statistical inference, we introduce an auxiliary variable $\delta \in \mathbb{R}^{P\times C}$,

$$\delta \sim \mathcal{MN}_{P\times C}(\delta_0, \Sigma_1, I_C) \quad \Rightarrow \quad \text{vec}(\delta) \sim \mathcal{N}_{PC}(\text{vec}(\delta_0), I_C \otimes \Sigma_1).$$

We define,

$$\text{vec}(\mathbf{B}) = (\Sigma_2^{1/2} \otimes I_P)\,\text{vec}(\delta) \tag{2}$$

vec($\mathbf{B}$) is a linear transformation of vec($\delta$) and thus follows a MNN

$$\text{vec}(\mathbf{B}) \sim \mathcal{N}_{PC}\left((\Sigma_2^{1/2} \otimes I_P)\,\text{vec}(\delta_0),\ (\Sigma_2^{1/2} \otimes I_P)(I_C \otimes \Sigma_1)(\Sigma_2^{1/2} \otimes I_P)^\top\right)$$

$$\text{vec}(\mathbf{B}) \sim \mathcal{N}_{PC}\left((\Sigma_2^{1/2} \otimes I_P)\,\text{vec}(\delta_0),\ \Sigma_2 \otimes \Sigma_1\right)$$

Substituting Eq. (2) into Eq. (1) result,

$$\mathbf{Y} = \mathbf{X}(\Sigma_2^{1/2} \otimes \mathbf{I}_P)\text{vec}(\delta) + \mathbf{E}, \mathbf{E} \sim \mathcal{N}(0, \sigma^2 I)$$

We further define $\mathbf{g} = \text{vec}(\delta)$, $\mathbf{g_0} = \text{vec}(\delta_0)$ and $\tilde{\mathbf{X}} = \mathbf{X}\left(\Sigma_2^{1/2} \otimes I_P\right)$. Eq. (1) changes to

$$\mathbf{Y} = \tilde{\mathbf{X}}\,\mathbf{g} + \mathbf{E} \tag{3}$$

### Prior distribution for g and $\sigma^2$

The conjugate priors are placed on $\mathbf{g}$ and $\sigma^2$

$$\mathbf{g}|\sigma^2 \sim \mathcal{N}(\mathbf{g}_0, \sigma^2(I_C \otimes \Sigma_1)), \quad \sigma^2 \sim \text{Inv-Gamma}(\alpha_0/2, b_0/2)$$

**Prior for $\mathbf{g}|\sigma^2$,**

$$p(\mathbf{g}|\sigma^2) \propto (\sigma^2)^{-CP/2} \exp\left(-\frac{1}{2\sigma^2}(\mathbf{g} - \mathbf{g}_0)^\top (I_C \otimes \Sigma_1)^{-1}(\mathbf{g} - \mathbf{g}_0)\right) \tag{4}$$

**Prior for $\sigma^2$,**

$$p(\sigma^2) = \frac{(b_0/2)^{\alpha_0/2}}{\Gamma(\alpha_0/2)}(\sigma^2)^{-(\alpha_0/2+1)} \exp\left(-\frac{b_0}{2\sigma^2}\right) \tag{5}$$

### Posterior distribution of g and $\sigma^2$

The **Likelihood** is

$$p(\mathbf{Y}|\mathbf{g}, \sigma^2, \tilde{\mathbf{X}}) \propto (\sigma^2)^{-N/2} \exp\left(-\frac{1}{2\sigma^2}\|\mathbf{Y} - \tilde{\mathbf{X}}\mathbf{g}\|^2\right) \tag{6}$$

Combining the prior from $\mathbf{g}$ (Eq. (4)), prior from $\sigma^2$ (Eq. (5)) and the likelihood (Eq. (6)), the joint posterior distribution of $\mathbf{g}$ and $\sigma^2$ is

$$\begin{aligned}
p(\mathbf{g}, \sigma^2|\mathbf{Y}, \tilde{\mathbf{X}}) &\propto p(\mathbf{Y}|\mathbf{g}, \sigma^2, \tilde{\mathbf{X}}) \cdot p(\mathbf{g}|\sigma^2) \cdot p(\sigma^2) \\
&\propto (\sigma^2)^{-N/2} \exp\left(-\frac{1}{2\sigma^2}\|\mathbf{Y} - \tilde{\mathbf{X}}\mathbf{g}\|^2\right) \\
&\quad \times (\sigma^2)^{-CP/2} \exp\left(-\frac{1}{2\sigma^2}(\mathbf{g} - \mathbf{g}_0)^\top (I_C \otimes \Sigma_1)^{-1}(\mathbf{g} - \mathbf{g}_0)\right) \\
&\quad \times \frac{(b_0/2)^{\alpha_0/2}}{\Gamma(\alpha_0/2)}(\sigma^2)^{-\alpha_0/2-1} \exp\left(-\frac{b_0}{2\sigma^2}\right).
\end{aligned} \tag{7}$$

We combine and simplify the exponential terms in Eq. (7)

$$\begin{aligned}
p(\mathbf{g}, \sigma^2|\mathbf{Y}, \tilde{\mathbf{X}}) &\propto (\sigma^2)^{-CP/2} \exp\left(-\frac{1}{2\sigma^2}\left[\mathbf{g}^\top\left((I_C \otimes \Sigma_1)^{-1} + \tilde{\mathbf{X}}^\top\tilde{\mathbf{X}}\right)\mathbf{g} - 2\mathbf{g}^\top\left((I_C \otimes \Sigma_1)^{-1}\mathbf{g}_0 + \tilde{\mathbf{X}}^\top\mathbf{Y}\right)\right]\right) \\
&\quad \times \left(\frac{1}{\sigma^2}\right)^{(\alpha_0+N)/2+1} \exp\left(-\frac{b_0 + \mathbf{Y}^\top\mathbf{Y} + \mathbf{g}_0^\top\left((I_C \otimes \Sigma_1)^{-1}\mathbf{g}_0\right)}{2\sigma^2}\right)
\end{aligned} \tag{8}$$

We now consider the quadratic form inside the first exponential of Eq. (8) and let

$$A = (I_C \otimes \Sigma_1)^{-1} + \tilde{\mathbf{X}}^\top\tilde{\mathbf{X}}, \quad \mu = (I_C \otimes \Sigma_1)^{-1}\mathbf{g}_0 + \tilde{\mathbf{X}}^\top\mathbf{Y}.$$

Based on this,

$$\mathbf{g}^\top A\mathbf{g} - 2\mathbf{g}^\top\mu + \mu^\top A^{-1}\mu - \mu^\top A^{-1}\mu = (\mathbf{g} - A^{-1}\mu)^\top A(\mathbf{g} - A^{-1}\mu) - \mu^\top A^{-1}\mu.$$

We rewrite the Eq. (8) as,

$$p(\mathbf{g},\sigma^2|\mathbf{Y},\tilde{\mathbf{X}}) \propto p(\mathbf{g}|\sigma^2,\mathbf{Y},\tilde{\mathbf{X}})p(\sigma^2|\mathbf{Y},\tilde{\mathbf{X}}) \tag{9}$$

$$= (\sigma^2)^{-CP/2}\exp\left(-\frac{1}{2\sigma^2}(\mathbf{g}-\mathbf{g}_n)^\top V_n^{-1}(\mathbf{g}-\mathbf{g}_n)\right)$$
$$\times (\frac{1}{\sigma^2})^{(\alpha_0+N)/2+1}\exp\left(-\frac{b_0+\mathbf{Y}^\top\mathbf{Y}+\mathbf{g}_0^\top\left((I_C\otimes\Sigma_1)^{-1}\mathbf{g}_0\right)-\mu^\top A^{-1}\mu}{2\sigma^2}\right) \tag{10}$$

**Posterior distribution of g**

The posterior distribution of $\mathbf{g}$ is a MVN

$$\mathbf{g}|\sigma^2,\mathbf{Y},\tilde{\mathbf{X}} \sim \mathcal{N}(\mathbf{g}_n,\sigma^2 V_n) \tag{11}$$

with posterior mean and variance of $\mathbf{g}$ as

$$\mathbf{g}_n = A^{-1}\mu = \left((I_C\otimes\Sigma_1)^{-1}+\tilde{\mathbf{X}}^\top\tilde{\mathbf{X}}\right)^{-1}\left((I_C\otimes\Sigma_1)^{-1}\mathbf{g}_0+\tilde{\mathbf{X}}^\top\mathbf{Y}\right) \tag{12}$$

$$V_n = A^{-1} = \left((I_C\otimes\Sigma_1)^{-1}+\tilde{\mathbf{X}}^\top\tilde{\mathbf{X}}\right)^{-1} \tag{13}$$

**Posterior distribution of $\sigma^2$**

The posterior distribution of $\sigma^2$ follows Normal-Inverse Gamma distribution

$$\sigma^2|\mathbf{Y},\tilde{\mathbf{X}} \sim \text{Inv-Gamma}(\alpha_n/2, b_n/2) \tag{14}$$

which includes an inverse gamma kernel

$$\left(\frac{1}{\sigma^2}\right)^{(\alpha_0+N)/2+1}\exp\left(-\frac{b_0+\mathbf{Y}^\top\mathbf{Y}+\mathbf{g}_0^\top(I_C\otimes\Sigma_1)^{-1}\mathbf{g}_0-\boldsymbol{\mu}^\top A^{-1}\boldsymbol{\mu}}{2\sigma^2}\right) \tag{15}$$

According to $\boldsymbol{\mu}^\top A^{-1}\boldsymbol{\mu} = \mathbf{g}_n^\top V_n^{-1}\mathbf{g}_n$,

Shape parameter:

$$\alpha_n = \alpha_0 + N \tag{16}$$

Scale parameter:

$$b_n = b_0 + \mathbf{Y}^\top \mathbf{Y} + \mathbf{g}_0^\top \left(I_C \otimes \Sigma_1\right)^{-1} \mathbf{g}_0 - \mathbf{g}_n^\top V_n^{-1} \mathbf{g}_n \tag{17}$$

**Posterior Distribution of vec(B)**

According to Eq. (2) and Eq. (11), the posterior distribution for vec($\mathbf{B}$) follows a MVN

$$\text{vec}(\mathbf{B})|\sigma^2, \mathbf{Y}, \tilde{\mathbf{X}} \sim \mathcal{N}\left((\Sigma_2^{1/2} \otimes I_P)\mathbf{g}_n, \sigma^2(\Sigma_2^{1/2} \otimes I_P)V_n(\Sigma_2^{1/2} \otimes I_P)^\top\right) \tag{18}$$

Therefore,

$$\mathbb{E}[\text{vec}(\mathbf{B})] = (\Sigma_2^{1/2} \otimes I_P)\mathbf{g}_n \tag{19}$$

$$\text{Cov}(\text{vec}(\mathbf{B})) = (\Sigma_2^{1/2} \otimes I_P)(\sigma_n^2 V_n)(\Sigma_2^{1/2} \otimes I_P)^\top \tag{20}$$

where $\sigma_n^2 = \mathbb{E}[\sigma^2] = \frac{b_n}{\alpha_n - 2}$ is the posterior expectation of $\sigma^2$ under the Inverse Gamma distribution, for $\alpha_n > 2$.

### Supplementary Figures

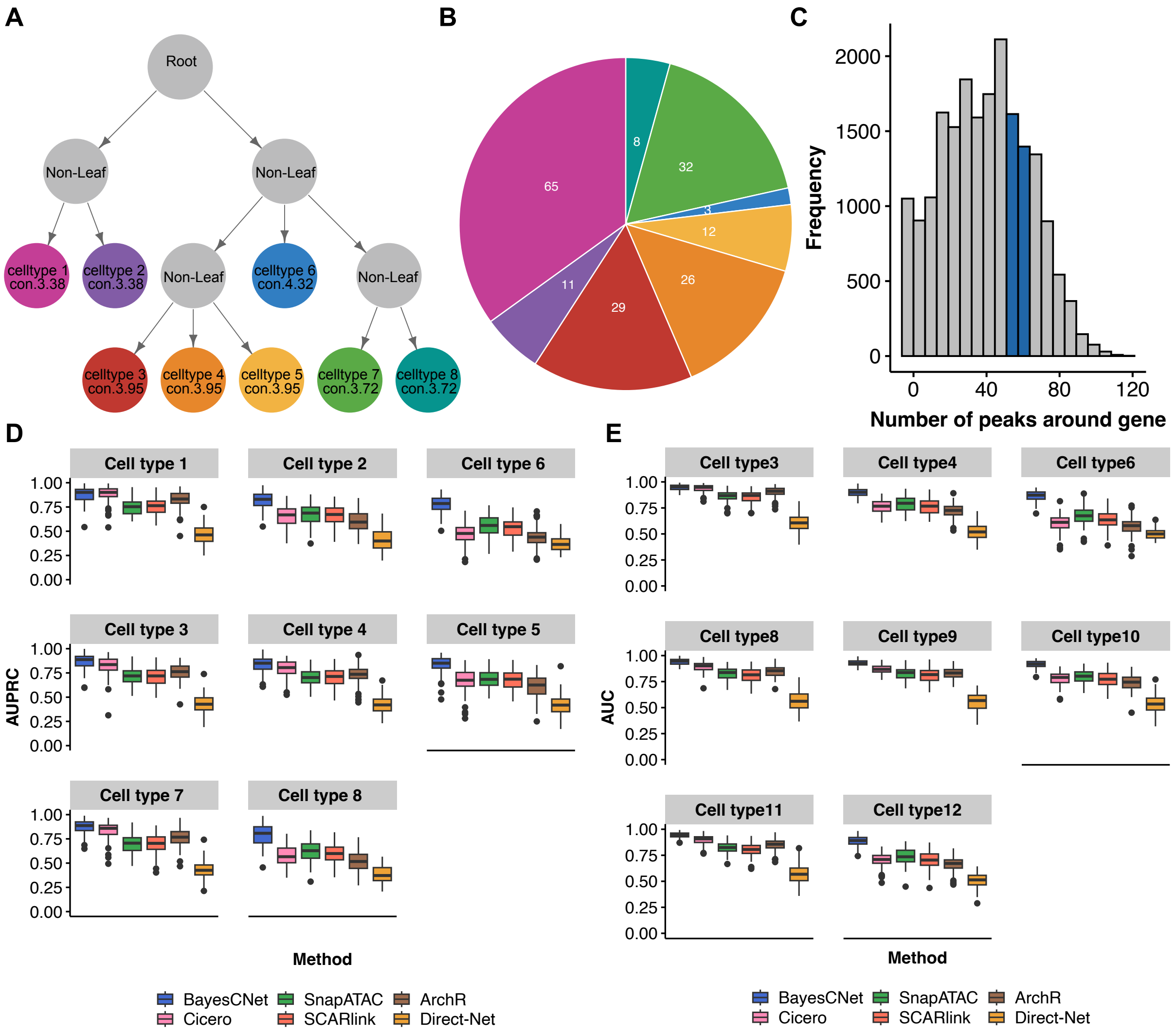


**Supplementary Figure 1. Real data simulation based on PBMC scMultiome dataset a.** A simulated cell ontology tree, mimicking PBMC cell type hierarchy, for simulation. **b.** Pie chart showing the distribution of metacells for each cell type in PBMC scMutiome dataset **c.** Histogram showing the distribution of the number of peaks within ±250 kb windows around each gene's transcription start site (TSS) in PBMC scMutiome dataset. Blue bar indicates the median number used in simulation. **d.** AUPRC across 100 simulations per method per cell type **e.** AUROC across 100 simulations per method per cell type

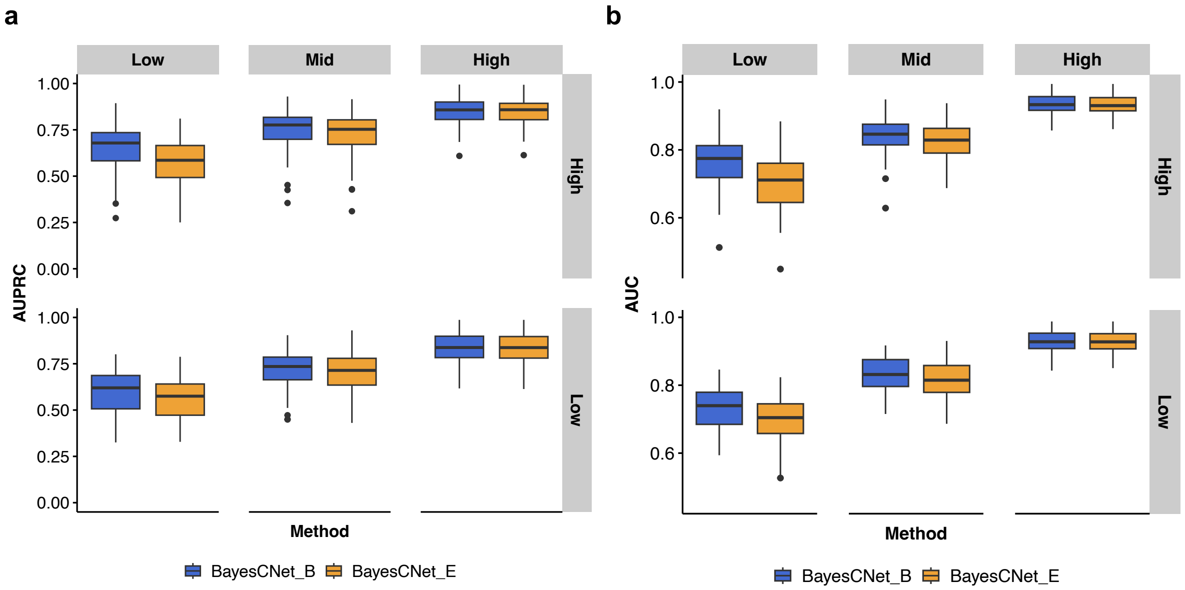


**Supplementary Figure 2. Simulation studies for comparing BayesCNet using biological prior (BayesCNet_B) inferred from PBMC cell ontology tree and empirical prior (BayesCNet_E) for inferring enhancer-gene linkages.** Two target cell types were selected to represent different connectivity levels: Cell type 2 (low connectivity) and Cell type 6 (high connectivity). Connectivity scores are indicated at each node. b. Experimental design of the 2×3 factorial simulation for connectivity and sample size. Each target cell type was simulated at three sample size levels: low (5), medium (20), and high (60) metacells, and two connectivity level: low and high. The sample sizes of other cell types are fixed at the medium level. **a.** AUPRC across 100 simulations per method per scenario. **b,** AUROC across 100 simulations per method per scenario.

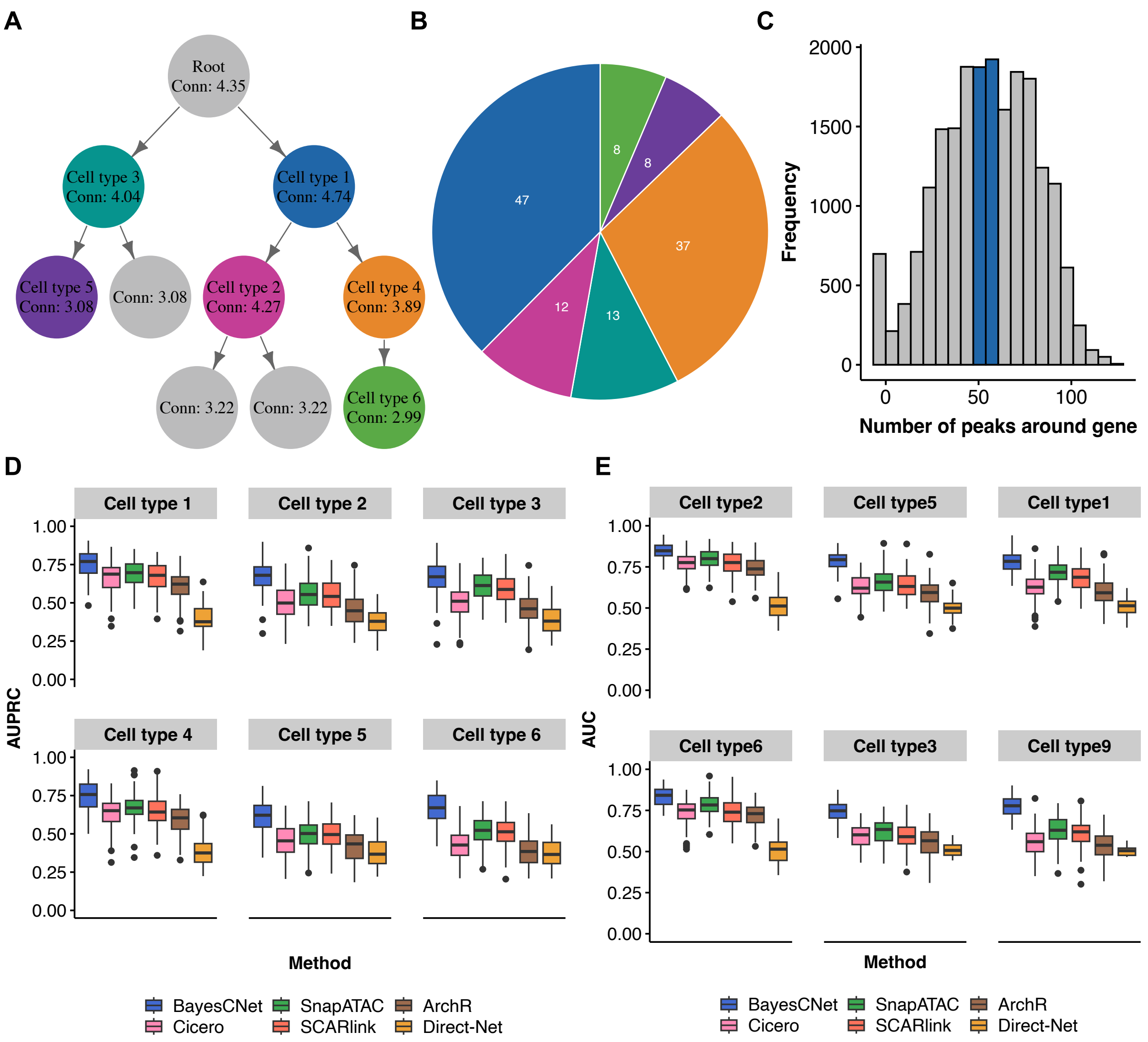


**Supplementary Figure 3. Real data simulation based on HEMA scMultimodal dataset** **a.** A simulated cell lineage tree, mimicking hematopoietic differentiation, for simulation. **b.** Pie chart showing the distribution of metacells for each cell type in HEMA scMultimodal dataset **c.** Histogram showing the distribution of the number of peaks within ±250 kb windows around each gene's transcription start site (TSS). Blue bar indicates the median number used in simulation. **d.** AUPRC across 100 simulations per method per cell type **e.** AUROC across 100 simulations per method per cell type

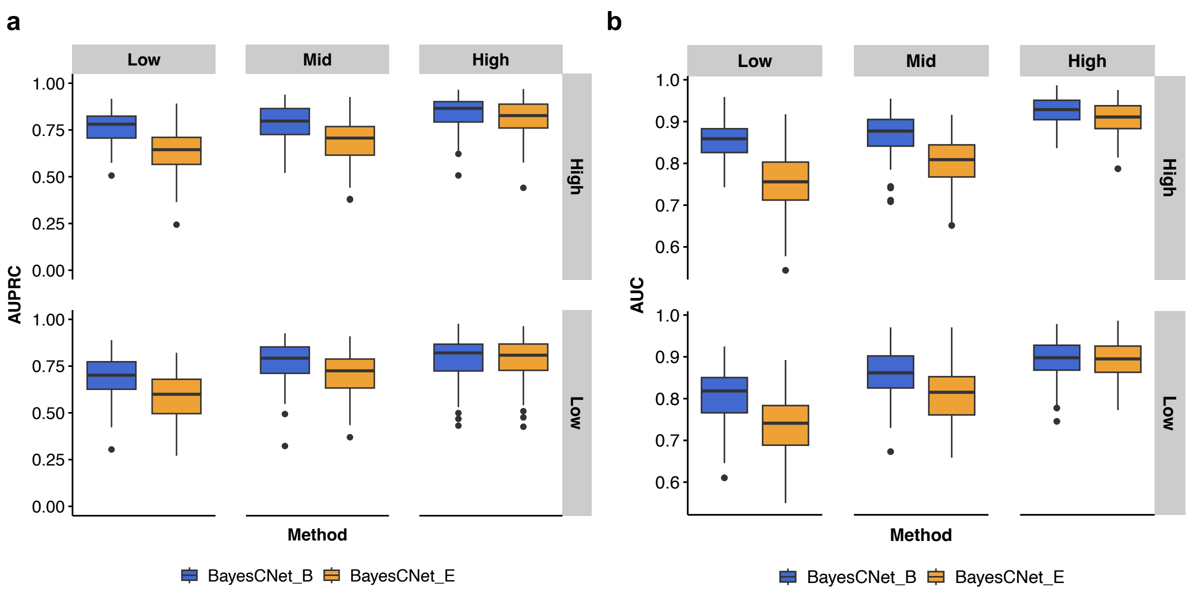


**Supplementary Figure 4. Simulation studies for comparing BayesCNet using biological prior (BayesCNet_B) inferred from HEMA cell lineage tree and empirical prior (BayesCNet_E) for inferring enhancer-gene linkages.** . Two target cell types were selected to represent different connectivity levels: Cell type 8 (low connectivity) and Cell type 3 (high connectivity). Connectivity scores are indicated at each node. b. Experimental design of the 2×3 factorial simulation for connectivity and sample size. Each target cell type was simulated at three sample size levels: low (8), medium (15), and high (40) metacells, and two connectivity level: low and high. The sample sizes of other cell types are fixed at the medium level. **a.** AUPRC across 100 simulations per method per scenario. **b,** AUROC across 100 simulations per method per scenario.

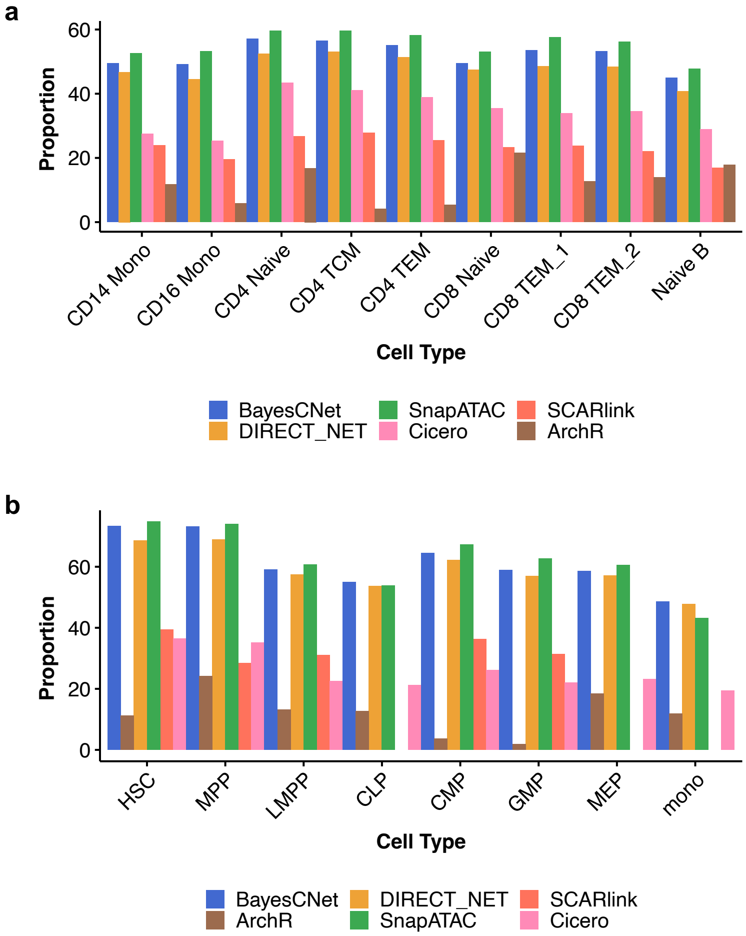


**Supplementary Figure 5. Proportion of evaluable cell type–specific marker genes in the PCHi-C analyses**. A marker gene was considered evaluable for a given method if at least five of its predicted EG pairs overlapped PCHi-C loop anchors in the matched cell type. **a.** Nine cell types from PBMC scMultiome dataset. **b.** Eight cell types from HEMA scMultimodal dataset.

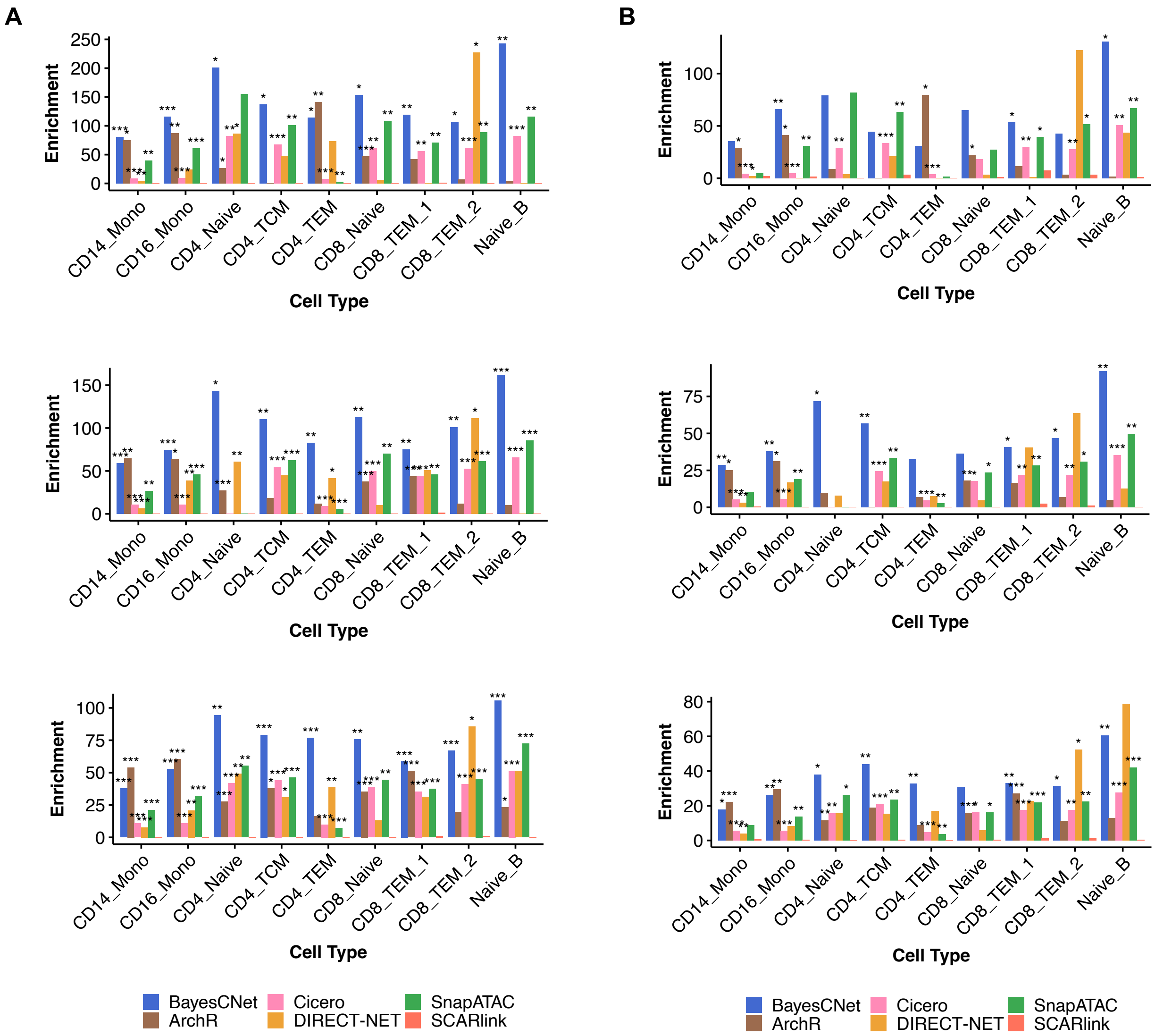


**Supplementary Figure 6 | Evaluating biological relevance of cell type-specific enhancer-gene linkages in PBMC scMutiome dataset using stratified linkage disequilibrium score regression (S-LDSC). a.** S-LDSC enrichment for systemic lupus erythematosus (SLE) (GWAS Catalog ID: GCST011097) using the enhancers from 10%, 25%, and 50% of predicted enhancer–gene linkages identified by each method across cell types in PBMC scMutiome dataset **b.** S-LDSC enrichment for SLE (GWAS Catalog ID: GCST005831) using the enhancers from 10%, 25%, and 50% of predicted enhancer–gene linkages identified by each method across cell types in PBMC scMutiome dataset. Asterisks indicate statistical significance ($p < 0.05$; $p < 0.01$; $p < 0.001$; $p < 0.0001$).

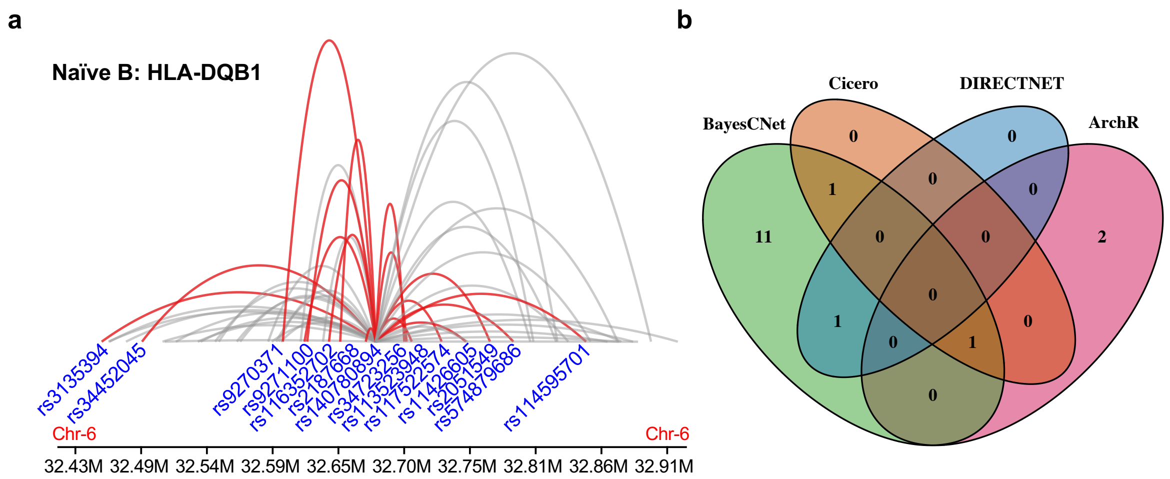


**Supplementary Figure 7. Colocalization between SLE-associated SNPs and *HLA-DQB1*.** **a** BayesCNet-identified Naïve B-specific enhancer–gene linkages colocalize with 14 SLE-associated SNPs at *HLA-DQB1* locus. All arcs denote BayesCNet-identified EG linkages. EG linkages overlapped with SLE-associated SNPs are highlighted in red. **b.** Venn diagram comparing SLE-associated SNPs in enhancers from EG linkages identified by all methods at the *HLA-DQB1* locus.

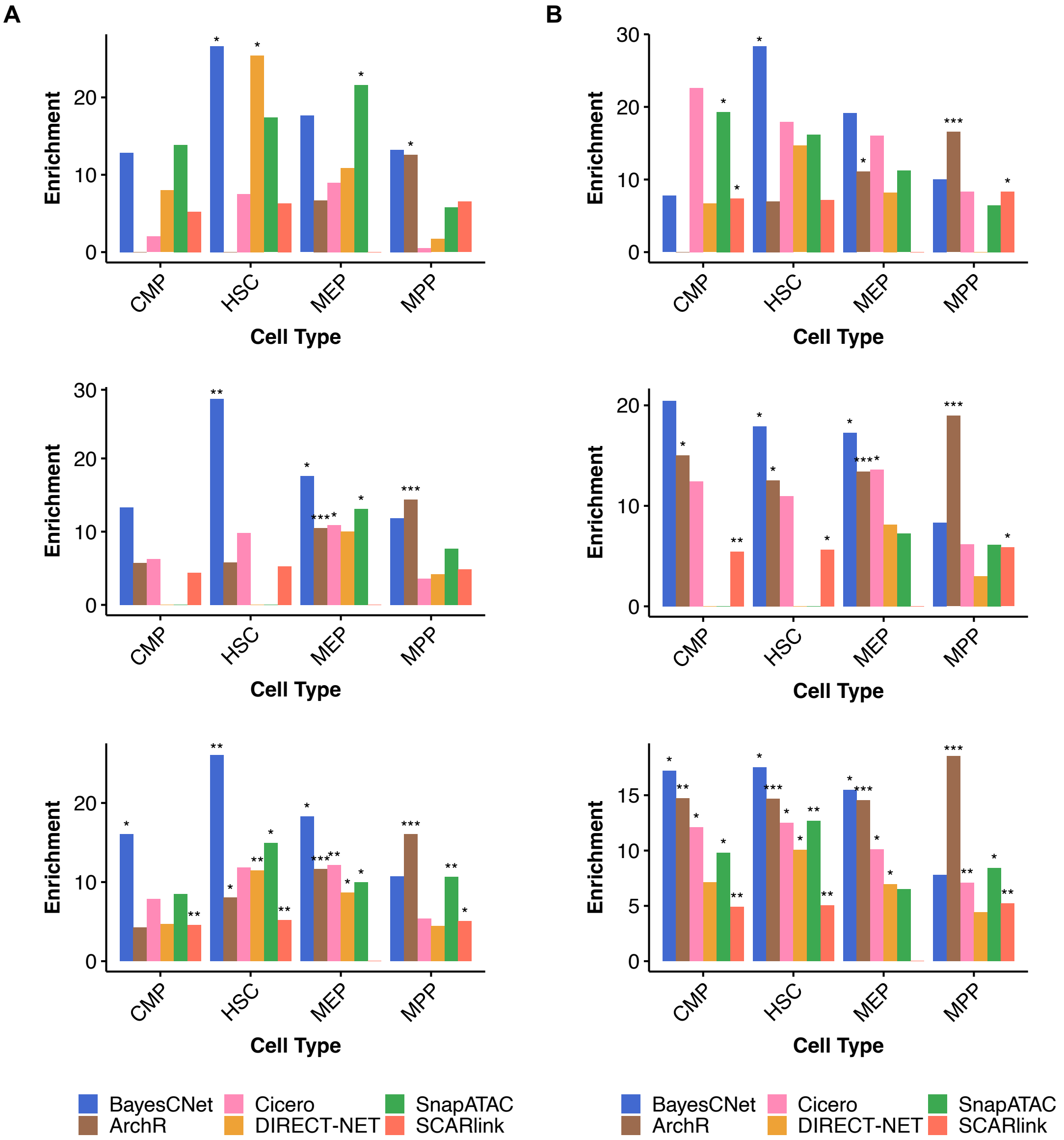


**Supplementary Figure 8. Evaluating biological relevance of cell type-specific enhancer-gene linkages using stratified linkage disequilibrium score regression (S-LDSC).** **a.**S-LDSC enrichment for platelet volume (GWAS Catalog ID: GCST90028996) based on enhancers from the top 10%, 25%, and 50% of predicted enhancer–gene linkages identified by each method across cell types in HEMA scMultimodal dataset. **b.**S-LDSC enrichment for platelet count (GWAS Catalog ID: GCST90468095) based on enhancers from the top 10%, 25%, and 50% of predicted enhancer–gene linkages identified by each method across cell types in HEMA scMultimodal dataset. Asterisks indicate statistical significance ($p < 0.05$; $p < 0.01$; $p < 0.001$; $p < 0.0001$).

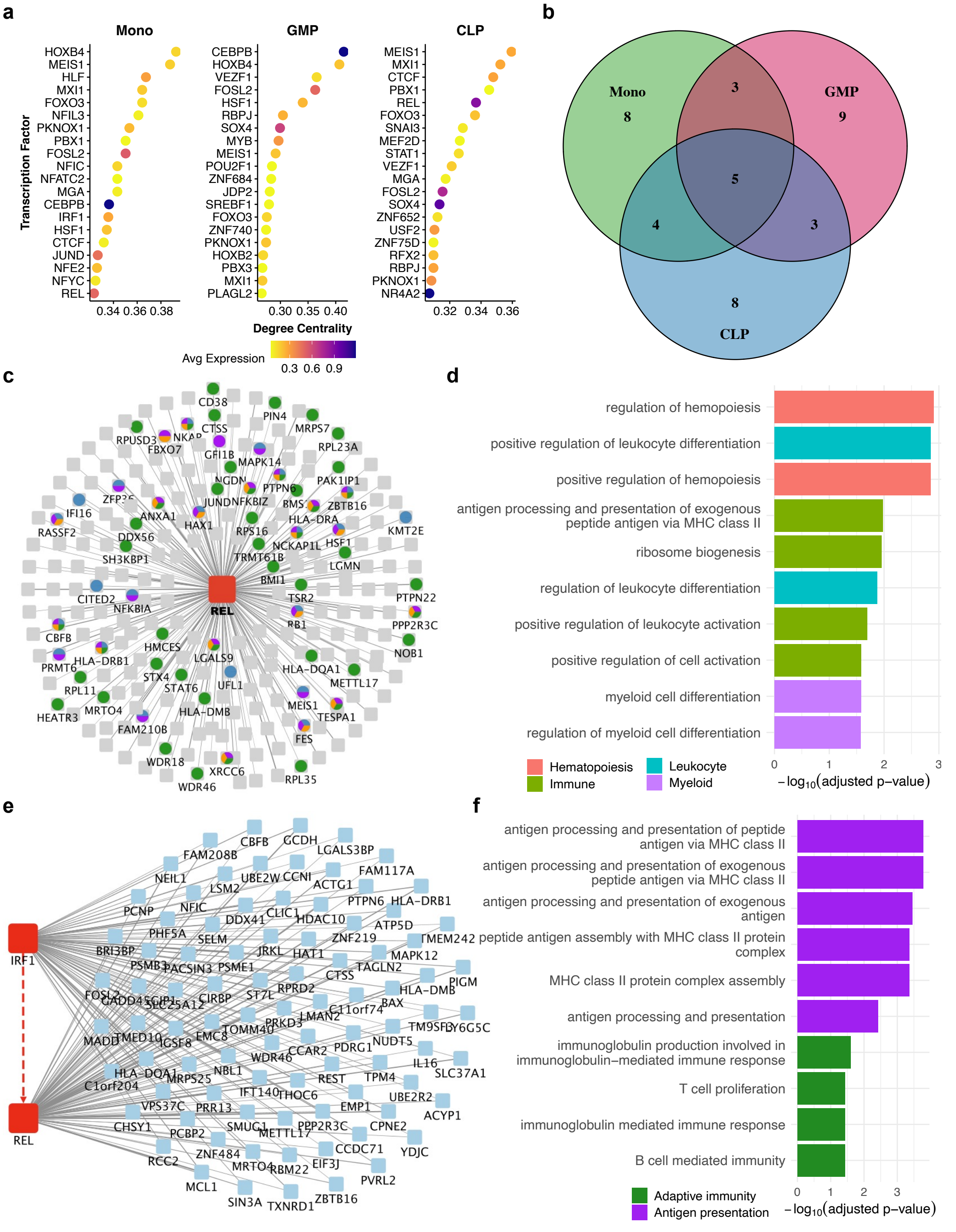


**Supplementary Figure 9. Cell type–specific TF–gene regulatory network (GRN) analysis in HEMA scMutimodal dataset using BayesCNet**. **a.**Top 20 transcription factors (TFs) ranked by degree centrality in Mono, GMP, and CLP. Node color indicates average TF expression. **b.** Venn diagram comparing top-ranked 20 TFs across the three cell types **c.** *REL*-hub TF-GRN in Mono **d.** Gene Ontology (GO) enrichment analysis of *REL* target genes **e.** Two interacted TF-GRN between *REL* and *IRF1* TF hubs in Mono **f.** GO enrichment of *REL* and *IRF1* shared gene targets.

### Supplementary Tables

**Supplementary Table 1.** Cell types, sample size of cells and metacells in PBMC scMultiome dataset.

| Cell Type | Single cell Count | Metacell Count |
|---|---|---|
| HSPC | 23 | 1 |
| CD14 Mono | 2790 | 65 |
| CD16 Mono | 521 | 11 |
| CD4 Naïve | 1332 | 29 |
| CD4 TCM | 1180 | 26 |
| CD4 TEM | 542 | 12 |
| Treg | 182 | 5 |
| CD8 Naïve | 1466 | 32 |
| CD8 TEM_1 | 302 | 8 |
| CD8 TEM_2 | 363 | 8 |
| MAIT | 108 | 3 |
| gdT | 107 | 2 |
| Naive B | 145 | 3 |
| Intermediate B | 357 | 8 |
| Memory B | 368 | 9 |
| Plasma | 18 | 1 |
| pDC | 85 | 2 |
| cDC | 198 | 5 |
| NK | 445 | 11 |

**Supplementary Table 2.** Cell types, sample size of cells and metacells in HEMA scMultimodal dataset.

| Cell Type | Full Name | Single cell Count | Metacell Count |
|---|---|---|---|
| HSC | Hematopoietic Stem Cell | 347 | 35 |
| MPP | Multipotent Progenitor | 142 | 15 |
| LMPP | Lymphoid-primed Multipotent Progenitor | 160 | 13 |
| CMP | Common Myeloid Progenitor | 502 | 47 |
| GMP | Granulocyte–Macrophage Progenitor | 402 | 37 |
| MEP | Megakaryocyte–Erythroid Progenitor | 138 | 12 |
| CLP | Common Lymphoid Progenitor | 78 | 8 |
| pDC | Plasmacytoid Dendritic Cell | 141 | 14 |
| Mono | Monocyte | 64 | 8 |

**Supplementary Table 3.** Cell types of the PCHi-C datasets.

| **Cell Type** |
|---|
| Megakaryocytes |
| Erythroblasts |
| Neutrophils |
| Monocytes |
| Macrophages M0 |
| Macrophages M1 |
| Macrophages M2 |
| Endothelial precursors |
| Naive B cells |
| Total B cells |
| Fetal thymus |
| Naive $CD4^+$ T cells |
| Total $CD4^+$ T cells |
| Non-activated total $CD4^+$ T cells |
| Activated total $CD4^+$ T cells |
| Naive $CD8^+$ T cells |
| Total $CD8^+$ T cells |

**Supplementary Table 4.** Matched cell type pairs between PBMC scMultiome datasets and PCHi-C datasets.

| **PBMC Cell Type** | **Matched PCHi-C Cell Type** |
|---|---|
| CD14 monocytes | Mono |
| CD16 monocytes | Mono |
| CD4 Naïve | Naive $CD4^+$ T cells |
| CD4 TCM | Total $CD4^+$ T cells |
| CD4 TEM | Total $CD4^+$ T cells |
| CD8 Naïve | Naive $CD8^+$ T cells |
| CD8 TEM_1 | Total $CD8^+$ T cells |
| CD8 TEM_2 | Total $CD8^+$ T cells |
| Naive B | Naive B cells |

**Supplementary Table 5.** Matched cell type pairs between HEMA scMultimodal datasets and PCHi-C datasets.

| **HEMA Cell Type** | **Matched PCHi-C Cell Type(s)** |
|---|---|
| HSC | All (exclude Endothelial precursors and Fetal thymus) |
| MPP | All (exclude Endothelial precursors and Fetal thymus) |
| LMPP | All T cells and B Cells |
| CLP | All T cells and B Cells |
| CMP | Monocytes, Macrophages, Neutrophil, Megakaryocyte, Erythroid |
| GMP | Monocytes, Macrophages, Neutrophil |
| MEP | Megakaryocyte, Erythroid |
| Mono | Monocytes |
